# Tumor-educated monocytes suppress T cells *via* adenosine and depletion of adenosine in the tumor microenvironment with adenosine deaminase enzyme promotes response to immunotherapy

**DOI:** 10.1101/2022.06.22.497260

**Authors:** Omar S. Sarkar, Howard Donninger, Numan Al Rayyan, Lewis C. Chew, Bryce Stamp, Xiang Zhang, Aaron Whitt, Chi Li, Melissa Hall, Robert A. Mitchell, Alfred Zippelius, John Eaton, Jason A. Chesney, Kavitha Yaddanapudi

## Abstract

Although immune checkpoint inhibitor (ICI) therapy has provided robust results in many cancer types such as melanoma and lung cancer, a large percentage of patients remain unresponsive to this therapy. Emerging evidence strongly suggests that one of the contributing factors in ICI resistance is monocytic myeloid derived suppressor cells (M-MDSCs) that accumulate in late-stage cancer patients. These M-MDSCs are a subset of innate immune cells and possess potent immunosuppressive activity against T lymphocytes. Here we provide evidence of a mechanism by which CD73-expressing M-MDSCs in the tumor microenvironment (TME) exhibit superior T cell suppressor function *via* adenosine. We show that tumor-derived PGE_2_, a prostaglandin frequently found at high levels in the TME, directly induces CD73 expression in M-MDSCs by initiating a signaling pathway that is mediated by both Stat3 and CREB. The resulting CD73 overexpression induces elevated levels of adenosine, a nucleoside with strong T cell suppressive activity, culminating in the suppression of CD8^+^ T cell-mediated anti-tumor responses. We also show that depletion of adenosine in the TME by the repurposed drug PEGylated Adenosine Deaminase (PEG-ADA) increases CD8^+^ T cell anti-tumor activity and enhances response to ICI therapy in preclinical models of cancer. Our results suggest that use of PEG-ADA is a viable therapeutic option to overcome ICI therapeutic resistance in advanced cancer patients.

## Introduction

The development of immune checkpoint inhibitors (ICIs; e.g., anti-PD-1 antibody) has revolutionized the therapy of several malignancies including melanoma, non-small cell lung cancer, and triple-negative breast cancer (1, 2). Despite the clinical success of ICIs in cancer treatment, therapeutic resistance continues to occur in large percentages of patients. One of the best predictors of clinical responses to anti-PD-1 antibody, is the presence of increased tumor somatic mutations (3). Relative mutational burden dictates neoantigen load that, in turn, elicits anti-tumor responses in the primary targets of anti-PD-1, the CD8^+^ T cells (4, 5). However, even among patients with high tumor neo-antigens and intrinsic numbers of antigen (Ag)-specific CD8^+^ T cells, up to 50% remain unresponsive to anti-PD-1 (5, 6). These findings strongly suggest the presence of additional mechanisms of immune suppression that directly confer resistance to ICIs and supports combinatorial treatment strategies to improve systemic disease control and patient survivability.

Several mechanisms have been proposed to account for the immune suppression in tumor-bearing hosts. One widely studied process involves cancer-induced aberrations in myelopoietic output, which lead to the generation and accumulation of different immunosuppressive myeloid cells in the tumor microenvironment (TME) of many solid tumors. These tumor-educated myeloid-derived suppressor cells (MDSCs) actively inhibit CD8^+^ T cell tumor homing and activation (7, 8). We and others have demonstrated that amongst the tumor-induced MDSC subsets, monocytic MDSCs (M-MDSCs) possess the greatest suppressive activity on CD8^+^ T cells (9, 10). Given that M-MDSC levels are elevated in human cancers (11, 12) and correlate with decreased patient survival (13, 14), intervention strategies that can reverse M-MDSC suppressive activity will support immunotherapy and enhance the durability of clinical response. Currently, such strategies are limited.

A further process that leads to immune suppression in tumors is mediated by the purine nucleoside Adenosine. Adenosine suppresses immune activity of T cells both directly by inhibiting activation of anti-tumor effector responses and indirectly by inducing expansion of other immune suppressive cell subtypes within the tumor microenvironment (15, 16). Metabolic changes occurring within a growing tumor favor the increased accumulation of extracellular adenosine (15). The generation of this purine nucleoside occurs through the degradation of ATP by the combined actions of enzymes CD39 (ectonucleoside triphosphate diphosphohydrolase; ATP → AMP) and CD73 (ecto-5′-nucleotidase; AMP → adenosine) (17). The biological rationale for this adenosine accumulation in normal animals is to limit the tissue damage in areas of injury. Cell damage arising from an injury leads to the release of intracellular ATP which is then converted to the anti-inflammatory compound adenosine. The latter, sensed by adenosine A2A receptors (18, 19) expressed on the immune cells, has the net effect of putting the brakes on inflammatory responses; however, in the case of cancer, the same activity suppresses immunity and facilitates tumor growth and metastasis. Identification of therapeutics that can effectively eliminate adenosine hold promise for disengaging the adenosine-mediated immune suppression in tumors.

Here, we show that prostaglandin E_2_ (PGE_2_) released from tumor cells directly induces expression of CD73 in M-MDSCs *via* a novel PGE_2_ → cAMP → CREB/STAT3-dependent pathway. We examine the phenotypic and functional behavior of CD73-expressing M-MDSCs in the tumor microenvironment (TME) in human lung cancer and in several types of murine cancers. We show that PGE_2_-induced CD73 sustains the immunosuppressive activity of M-MDSCs in the TME. We further show that CD73-expressing M-MDSCs exhibit superior T cell suppressor function *via* an adenosine-mediated mechanism.

In order to evaluate if therapeutically targeting the immunosuppressive intratumoral adenosine can improve the efficacy of ICI therapy, we use a novel agent, <u>P</u>oly<u>e</u>thylene <u>G</u>lycol-conjugated [PEGylated]-Adenosine Deaminase (PEG-ADA) (20). PEG-ADA is FDA-approved as an enzyme replacement therapy for children with severe combined immunodeficiency (SCID) (20). This enzyme converts adenosine to the inert, non-immune suppressive metabolite, inosine. Our data indicate that depletion of adenosine by PEG-ADA relieves adenosine-mediated T cell immune suppression and sensitizes tumors to ICI therapy.

## Materials and Methods

### Patient Samples

Deidentified tumor and adjacent normal tissues, and peripheral blood were collected from 19 Stage II/III non-small cell lung cancer (NSCLC) patients. NSCLC patients included in this study were not undergoing therapy when their samples were collected. Patient samples were collected after informed consent was obtained by staff of the UofL Health-Cancer Center Biorepository and covered under University of Louisville IRB protocol number 08.0388. Fresh tumor tissues were processed into single-cell suspensions for phenotype and functional studies or flash frozen for adenosine measurements.

### Cell lines

The human melanoma cell line A375 (ATCC CRL-1619), human lung cancer line A549 (ATCC CCL-185), mouse melanoma line B16-F10 (ATCC CRL 6475), mouse lung cancer line LLC (ATCC CRL-1642), and mouse breast cancer cell line 4T1 (ATCC CRL-2539), were purchased from ATCC (Manassas, VA) and maintained in DMEM (Corning) or RPMI 1640 (Corning) media containing 10% (v/v) FBS. KP1.9 cells were obtained from Dr. Zippelius’ lab. 4T1-sh*PTGES* and 4T1-shScr were generated by stable induction of lentiviral particles containing expression constructs for sh*PTGES* or Scr control, respectively (Origene Technologies) and selection in Puromycin. Cells were frozen at passage two after receiving them from ATCC and were maintained in culture no longer than 6–8 weeks. All cell lines were authenticated by the ATCC cell bank using the Short Tandem Repeat (STR) profiling.

### Biochemical reagents

Human and murine GM-CSF, IL-6, IL-4, and IL-10 were obtained from Peprotech. PGE_2_ and dibutyryl-cAMP were obtained from Tocris. The EP2 receptor agonist Butaprost and EP4 receptor agonist CAY10598 were obtained from Cayman. Forskolin was obtained from Enzo Life Sciences. STAT3 inhibitor S3I-201 and CREB inhibitor 666-15 were obtained from Calbiochem. Adenosine, ADA, AMP, AMP-CP, erythro-9-(2-hydroxy-3-nonyl) adenine (ADA inhibitor), ovalbumin, and human serum were obtained from Sigma-Aldrich. Neutralizing anti-IL-10 antibody was obtained from R&D systems. Carboxyfluorescein succinimidyl ester (CFSE) was obtained from Thermo Fisher Scientific.

### Mice

All mice were handled in accordance with the Association for Assessment and Accreditation of Laboratory Animals Care international guidelines, with the approval of the Institutional Animal Care and Use Committee at University of Louisville. The University of Louisville IACUC reviewed and approved this study under ID # 21972. Wild-type C57BL/6J and BALB/c, OT-I TCR transgenic, and Nt5e^-/-^ C57BL/6 mice were purchased from the Jackson Laboratory. Mice were bred and housed in a pathogen-free barrier animal facility and maintained on a standard 12-hour light/12-hour dark cycle.

### *In vivo* tumor experiments]

#### Implantable tumor models

Syngeneic male 6–8-week-old C57BL/6 mice were inoculated subcutaneously (s.c.) with 2 × 10^5^ LLC, B16-F10 or KP1.9 cells. Female 6–8-week-old Balb/c mice were inoculated s.c with 2 × 10^5^ 4T1-shScr control cells or 4T1-*shPTGES* cells in the left flank.

#### Orthotopic NSCLC model

Confluent KP1.9 cells were detached with trypsin to obtain single-cell suspensions. 0.5 × 10^6^ KP1.9 cells were injected in the tail vein of 8–10-week-old wild-type C57BL/6 or Nt5e^-/-^ C57BL/6 mice.

#### Tumor monitoring

(i) Subcutaneous tumor growth was monitored every 2 days using digital calipers to measure both the longitudinal and transverse diameters (in mm). Animals bearing tumors were euthanized when tumors reached a volume of 1500 mm^3^ or earlier if tumors ulcerated or animals showed signs of discomfort; (ii) Orthotopic lung tumor growth was monitored using micro-CT imaging using a MicroCAT II scanner (Siemens). Mice were anesthetized with isoflurane during the entire imaging procedure. Imaging was started at day 24 post-tumor inoculation and continued once a week until day 42; and (iii) For histological analysis of tumor-bearing lung tissues, tissue sections of the lungs were prepared as previously described, with minor modifications (21). Briefly, before organ collection, intracardiac perfusion was performed with ice-cold PBS and lung tissues were fixed in 10% neutral phosphate-buffered formalin for 24 hours at room temperature. After the tissues were embedded in paraffin blocks, paraffin microtomy (RM2135; Leica Biosystems) was performed at 5 μm thickness per section. To stain with hematoxylin (95057-844; VWR; Radnor, PA) and eosin (HT110232; Thermo Fisher), sections were deparaffinized and rehydrated in xylene, ethanol, and deionized water and stained (21). Stained slides were dehydrated with ethanol, which was then extracted with xylene. Xylene-based PermountTM mounting medium (SP15-500; Thermo Fisher; Waltham, MA) was used to affix the coverslips to slide. Slides were scanned using an Aperio Imagescope (Leica Biosystems).

#### Treatment of tumor-bearing mice

(i) PEG-ADA monotherapy – KP1.9 NSCLC cells were injected at 1 × 10^5^ cells per mouse s.c. Once the tumors reached 50 mm^3^ in size, two groups of KP1.9-bearing mice (n = 10 mice per group) received the following treatments every other day *via* the intraperitoneal route: 1) vehicle control (saline); 2) PEG-ADA (2 Units per mouse; procured from Leadiant Biosciences, Gaithersburg, Maryland, USA). Tumors were measured by digital calipers and plotted as volume/time; and (ii) PEG-ADA and anti-PD-1 combination therapy – Four groups of KP1.9-bearing mice (n = 8 mice per group) received the following treatments every other day *via* the intraperitoneal route: 1) vehicle control (saline), 2) anti-PD-1 Ab alone (clone RMPI-14, 250 μg/mouse, Bio X Cell), 3) PEG-ADA alone (2 Units per mouse), and 4) anti-PD-1 plus PEG-ADA.

End points for experiments with mice were selected in accordance with institutional-approved criteria. All tumor cell lines used in the *in vivo* experiments were routinely tested for mycoplasma contamination using Mycoalert plus mycoplasma detection kit (Lonza). Randomization of animals was not used in experiments and no blinding was done for the animal experiments.

### *In vitro* generation of human and murine M-MDSCs

(i) To generate tumor cell line-induced human M-MDSCs, CD14^+^ cells (1 × 10^6^) isolated from PBMCs obtained from healthy donors were co-cultured with A375 or A549 tumor cells (5 × 10^5^) in complete IMDM medium (IMDM + 10% human serum) per well in a 6-well transwell system separated by a 0.4 μm membrane. A375 and A549 co-cultured monocytes and control monocytes cultured without tumor cells were harvested from the bottom wells by gently scraping after 68–72 hours of culture; (ii) To generate PGE_2_-induced human M-MDSCs, CD14^+^ cells (1 × 10^6^) isolated from PBMCs obtained from healthy donors were treated with GM-CSF (10 ng/mL) and PGE_2_ (1 μM) for 6 days. Half of the culture medium was replaced with fresh medium containing GM-CSF/PGE_2_ every 48 hours. In some experiments, Butaprost (10 μM) and CAY10598 (10 nM) were added at the beginning of the 6-day culture; and (iii) To generate murine PGE_2_-induced M-MDSCs, fresh bone marrow (BM) cells were harvested from the tibia and femur of C57BL/6 mice using a previously established protocol (9). BM cells were cultured for 5 days in complete RPMI medium (RPMI + 10% FBS/ 10mM HEPES/20 μM BME) with murine GM-CSF (40 ng/mL) and PGE_2_ (2.5 μM). Half of the medium was replaced every 48 hours with fresh bone marrow medium containing GM-CSF/PGE_2_.

### *In vitro* generation of human and murine Dendritic cells (DCs)

(i) To generate human monocyte-derived DCs, CD14^+^ cells (1 × 10^6^) isolated from PBMCs obtained from healthy donors were treated with GM-CSF (10 ng/mL) and IL-4 (10 ng/mL) for 6 days. Half of the culture medium was replaced with fresh medium containing GM-CSF/IL-4 every 48 hours; and (ii) To generate murine DCs, BM cells were cultured for 5 days in complete RPMI medium with murine GM-CSF (40 ng/mL) and IL-4 (10 μM). Half of the medium was replaced every 48 h with fresh bone marrow medium containing GM-CSF/IL-4.

### Isolation of murine lung mononuclear cells (LMCs)

LMCs were isolated as described previously with modifications (22). Briefly, single-cell suspensions were prepared first by cutting lungs into ∼1-mm segments, followed by incubation at 37 °C for 45 minutes in RPMI 1640 with Liberase TM (50 μg/mL) and DNase I (1 μg/mL). An isotonic discontinuous Percoll (GE Healthcare) density gradient [30:70% (v/v)] was then used for mononuclear cell isolation.

### Monocytic MDSC isolation

(i) Human CD14^+^CD73^+^ tumor-associated monocytes were positively selected from human NSCLC lesions. Fresh tumor tissue from NSCLC patients were enzymatically digested and single-cell suspensions were stained with CD14 microbeads (Miltenyi Biotec) and positively selected using AutoMACS ProSeparator (Miltenyi Biotec). CD14^+^ cells were then stained with biotinylated anti-human CD73 antibody and streptavidin microbeads, and CD14^+^CD73^+^ cells were separated using AutoMACS ProSeparator; and (ii) Murine CD11b^+^Gr-1^int^ M-MDSCs were positively selected from PGE_2_-induced BM-MDSCs and from LMCs obtained from naïve, KP1.9-bearing wild-type and Nt5e ^-/-^ mice using the murine MDSC isolation kit (Miltenyi). For purifying CD73^+^ and CD73^neg^ murine M-MDSCs, purified M-MDSC cells were staining with biotinylated anti-mouse CD73 antibody and streptavidin microbeads and were sorted using AutoMACS Pro Separator.

### Flow Cytometry analysis

For surface staining, single cell suspensions were washed and resuspended in PBS, stained with Fixable Viability Dye eFluor™ 450 (from Thermo Fisher Scientific, USA) for 30 minutes. Cells were then washed twice and resuspended in FACS buffer [2% FBS + PBS] and stained with multi-color antibody (Ab) cocktails according to the manufacturer’s recommendations. Antibodies used in this study are provide in **Supplemental Table 1**. Murine cells were incubated with CD16/CD32 antibody (2.4G2) for 10 min at 4 °C to block Fc receptors before being stained for surface markers. For intracellular staining, cells were fixed in Fix/Perm solution (BD Biosciences). After being washed with Perm/ Wash buffer (BD Biosciences), the cells were stained intracellularly for 30 minutes, washed, and resuspended in flow buffer. Stained cells were analyzed with a FACSCanto II [Becton Dickinson (BD) Biosciences, Franklin Lakes, NJ] and data were analyzed using FlowJo v10 (BD Biosciences, Franklin Lakes, NJ). FACS dot plots are represented with log-scale axes. Histograms are represented on a log-scale x-axis and a linear y-axis.

### Immunosuppressive assays

The suppressive function of human and murine M-MDSCs was performed using our previously established protocol (9). For human M-MDSC suppressive assays, autologous T cells were isolated from patient NSCLC lesions or from healthy donor PBMCs using the Pan T-cell Isolation Kit (Miltenyi Biotec). Autologous T cells were co-cultured with either: 1) CD14^+^CD73^+^ cells from patient tumors, 2) A375/A549 cell line-induced M-MDSCs, 3) PGE_2_-induced M-MDSCs, 4) CD73^+^ and CD73^neg^ M-MDSCs, or 5) CD73^+^ MDSC–derived supernatants that were untreated or treated with ADA. Cultures were then stimulated with either anti-CD3 and anti-CD28 antibodies or tetanus toxoid (1.0 μg/mL) (in 96-well flat-bottomed plates) or anti-CD3/anti-CD28 beads (in 96-well round-bottomed plates) for 4 or 5 days. T cell proliferation was measured by ^[3H]^thymidine incorporation. For analysis of proliferation with flow cytometry, autologous T cells were pre-labelled with CSFE (5 μM); and (ii) For murine M-MDSC suppressive assays, splenocytes from OTI transgenic mice were co-cultured with M-MDSCs from BM or KP1.9 tumor-bearing lungs at different ratios in the presence of ovalbumin (250 μg/mL) for 4 days and proliferation of CD8^+^ T cells were measured. In some experiments, M-MDSCs were pretreated with CD73 the inhibitor AMP-CP (10 μM) and AMP for 8 hours and supernatants were added to T cells and proliferation was measured.

### Quantitative PCR analysis

RNA was extracted from CD14^+^ cells using the RNAeasy micro kit (Qiagen) following the manufacturer’s instructions. Reverse transcription was performed with the iScriptTM cDNA synthesis kit (Bio-Rad). Quantitative mRNA assessment was performed by real-time reverse transcription PCR using Taqman Universal Master Mix (Applied biosystem) as previously described (9, 23). The following TaqMan probes from Applied Biosystems were used according to manufacturer’s instructions: 18s (Hs99999901_s1; VIC), CD73 (Hs00159686_m1; FAM), CD39 (Hs00969559_m1), IL-6 (Hs00174131_m1; FAM), IL-10 (Hs00961619_m1; FAM), and A2AR (Hs00169123_m1; FAM).

### ELISA

PGE_2_ was measured by ELISA (R&D Systems) in supernatants from murine LLC, KP1.9, and B16-F10 cell cultures. IL-6 and IL-10 levels were measured by ELISA in supernatants from human CD14^+^ monocytes. Human IL-6 ELISA kit was obtained from Boster Bio and Human IL-10 ELISA kit was obtained from R&D Systems.

### Adenosine measurement

Adenosine in tumors from NSCLC patients and in M-MDSC culture supernatants was measured by liquid chromatography coupled with tandem mass spectrometry. Adenosine was measured in fresh culture supernatants of CD73-M-MDSCs treated with AMP (10 μM) for 6 hours and erythro-9-(2-hydroxy-3-nonyl) adenine (EHNA), an inhibitor of adenosine deaminase, was added to culture supernatant to prevent adenosine degradation over-time. 1 mg of tumor tissue was homogenized in 25μL of 80% MeOH. Gentle N_2_ flow was applied to evaporate the MeOH. Freeze-fried samples were reconstituted with 400 μL of water and 2 μL of the injection volume was used for LC-MS analysis. The linear range for adenosine quantification on LC/MS instrument was determined as 50 nM to 50,000 nM. Multiple reaction monitoring (MRM) method was applied to qualify and quantify adenosine.

### Immunoblotting

CD14^+^ cells were cultured in complete IMDM (IMDM + 10% human serum) and were treated with GM-CSF alone (10 ng/mL) or GM-CSF/PGE_2_ (1 μM), GM-CSF/IL-6 (10 ng/mL), Forskolin (10 μM), dibutyryl-cAMP (100 μM), STAT3 inhibitor (S3I-201; 100 μM), CREB inhibitor (666-15; 2 μM) and STAT3i/CREBi combination. Cells were lysed in RIPA buffer (Sigma) supplemented with a protease inhibitor cocktail (Sigma) and equal amounts of proteins analyzed by western blot using a Novex NuPage gel system (Thermofisher) using the following antibodies; from Cell Signaling Technology; Phospho-Stat3 (Tyr705; 9138), Stat3 (9139), Phospho-NF-κB (Ser536; 3033), NF-κB (8242), IκBα (9242), phospho-CREB (Ser133; 9198), and CREB (9197), and from Santa Cruz Biotechnology; PTGES (sc365844).

### Statistical Analysis

GraphPad Prism 8.0 software (GraphPad Prism Software, Inc., La Jolla, CA) and SAS version 9.4 (SAS Institute, Inc, Cary, NC) were used for statistical analyses. Two-group comparisons between control and test samples (groups compared are indicated in the respective figures) were done by two-tailed Student’s *t* tests. Multiple data comparisons were derived by one-way ANOVA followed by Tukey’s post-hoc test. Correlations between PGE_2_ levels and CD73^+^ M-MDSC frequencies were assessed using Spearman’s Rank Correlation. For all tests, statistical significance was assumed where *p<0*.*05*. For all figures, *, **, and *** denote p < 0.01, p < 0.001, and p < 0.0001, respectively.

## Results

Previously, we had demonstrated that the frequencies of CD14^+^HLA-DR^low/–^ M-MDSCs are significantly elevated in patients with advanced stage cancer and we established an *in vitro* model of human M-MDSC induction using healthy donor monocytes co-cultured with the melanoma cell line, A375 (9). A comparative microarray analysis of cultured monocytes and A375-educated monocytes (A375-MDSCs) revealed that several immunosuppressive mediators including IL-6, IDO, and NOX4 are differentially expressed in A375-M-MDSCs (9). One gene product of particular interest that was upregulated to high levels in A375-MDSCs is the ecto-5′-nucleotidases CD73 (**Supplementary Fig. 1A**). Given that CD73 is centrally involved in generating adenosine, a potent immunosuppressive molecule in the TME (24), we validated the high expression levels of *CD73* in A375-MDSC by qPCR (**Supplemental Fig. 1B**). Intriguingly, we found that the expression of *CD39*, an ecto-nucleoside triphosphate diphosphohydrolase enzyme that converts ATP to AMP, is downregulated in A375-MDSCs (**Supplemental Figs. 1A and C**).

To determine if soluble factors released from the tumor cells influence CD39/CD73 expression in M-MDSCs, we co-cultured A375 and A549 cells with CD14^+^ monocytes from healthy donors using an *in vitro* transwell experimental system; monocytes cultured in tumor cell-free media were used as the control. As shown in **Figs. 1A–C**, CD73 expression is significantly higher in lung cancer- and melanoma-educated CD14^+^HLA-DR^low^/^neg^ monocytes when compared to that in monocytes cultured in media alone. In monocytes exposed to tumor cell media, we observed significant increases in the percentage of CD73-bearing cells (**Figs. 1A and B**) as well as the relative expression levels of CD73 (**MFI; Fig. 1C**). The expression levels of CD39, on the other hand, were very low in both the cultured and tumor-educated monocytes (**Supplemental Fig. 1D**). Importantly, we found that these tumor-educated CD73^+^ monocytes significantly inhibited the proliferation of autologous T cells in an *in vitro* T cell suppressive assay (**Figs. 1D and E**).

**Figure 1.**
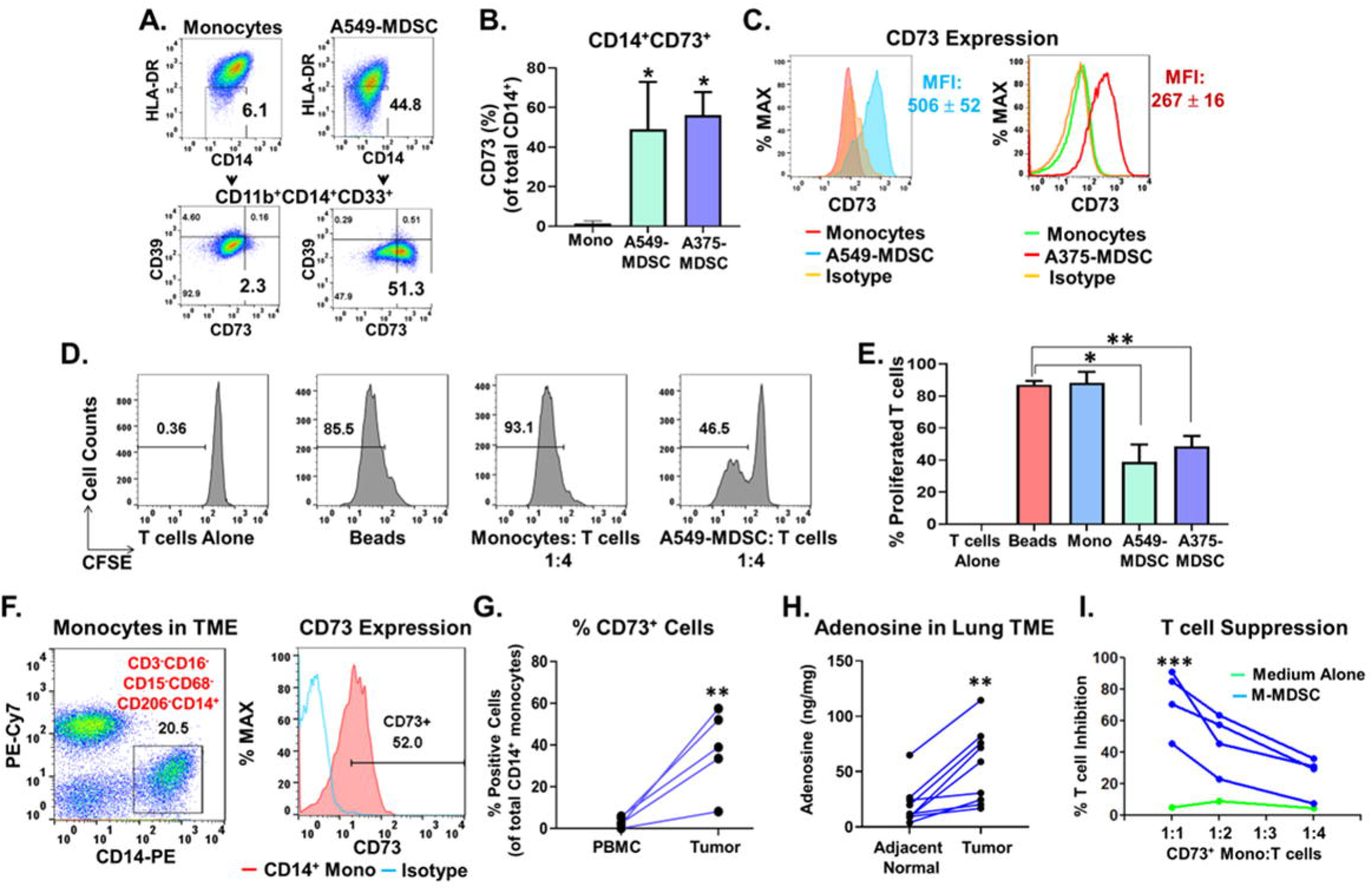
Tumor-associated immunosuppressive myeloid cells express CD73. **A–C**. Tumor cells induce CD73 expression in monocytic MDSCs *via* a soluble factor. CD14^+^ monocytes from healthy donor PBMCs were co-cultured with tumor cells using a transwell system. Representative dot plots **(A)**, bar graph **(B)**, and histograms **(C)** showing the expression of CD73 (proportions and MFI) in A549 and A375 tumor cell-induced human M-MDSCs. Data, mean ± SEM of three independent experiments with monocytes from 3 different donors, *P* values: *, p<0.01. **D, E**. Immune suppressive activity of A549- and A375-induced M-MDSCs. Autologous CFSE-labeled T cells were cultured in the presence of monocytes cultured for 64 hours in the absence of tumor cells (cultured mono), or with monocytes co-cultured with A549 or A375 cells (A549-MDSC and A375-MDSC). T cells were activated with anti-CD3/anti-CD28 beads in the absence or presence of the indicated monocytes/MDSCs for 4 days. Representative histograms **(D)** and bar graph **(E)** showing the percentage of proliferated CFSE-labeled T cells. Data, mean ± SEM of three independent experiments, *P* values: **, p<0.001; *, p<0.05. **F, G**. Human non-small lung cancer (NSCLC)-associated CD14^+^CD68^neg^CD206^neg^ monocytes present in the tumors express higher levels of CD73 when compared to expression levels in the circulating monocytes. Data, mean ± SEM of n = 4 patients, *P* values: **, p<0.001. **H**. Adenosine levels are consistently higher in NSCLC tumors when compared with surrounding healthy tissue. Tumor and adjacent healthy tissues were resected from the same NSCLC patients (from n = 9 patients) and adenosine levels were quantified using LC-MS. *P* values: **, p<0.001. **I**. Immunosuppression mediated by human NSCLC-associated CD73^+^ monocyte cells. CD3^neg^CD14^+^CD73^+^ monocyte cells and autologous CD3^+^ cells were sorted from fresh NSCLC cancer tissues and co-cultured for 4 days in the presence of anti-CD3/anti-CD28 antibody. Representative bar graph **(I)** showing the percentage of T cell inhibition. Data, mean ± SEM of cells from 4 different patients.

We extended these tumor cell line-based studies to human primary lung cancer. CD73 expression was significantly higher in metastatic lung tumor-infiltrating CD14^+^HLA-DR^+^CD68^neg^CD15^neg^CD16^neg^ monocytes when compared to CD73 expression in circulating CD14^+^ monocytes (**Figs. 1F and G**). Next, we assessed the contribution of CD73 enzymatic activity to adenosine generation and immune suppression in the lung TME. Analysis using Liquid Chromatography / Mass spectrometry (LC/MS) revealed higher levels of adenosine in lung tumor tissue than in adjacent normal tissue (**Fig. 1H and Supplemental Fig. 2A**). To characterize the functional phenotype of CD73-expressing CD14^+^ monocytes from human lung cancer samples, we sorted these cells and examined their suppressive activity. Tumor-infiltrating CD73-expressing CD14^+^ cells inhibited the CD3/anti-CD28-induced proliferation of autologous CD3^+^ T cells (**Fig. 1I**).

To determine which soluble factors from the tumor cells are responsible for the observed upregulation of CD73, we exposed the CD14^+^ monocytes isolated from healthy donors to previously reported suppressive factors released from tumor cells—granulocyte colony-stimulating factor (GM-CSF), PGE_2_, IL-6, and IL-4 or a combination of these factors. GM-CSF alone, or GM-CSF + IL-6 or GM-CSF + IL-4 combinations did not markedly alter the surface CD73 expression on monocytes (**Figs. 2A and B**). However, the short-term exposure of the monocytes to the GM-CSF + PGE_2_ combination resulted in significantly enhanced levels of CD73 expression than that exhibited in monocytes exposed to GM-CSF alone (**Figs. 2A and B**).

**Figure 2.**
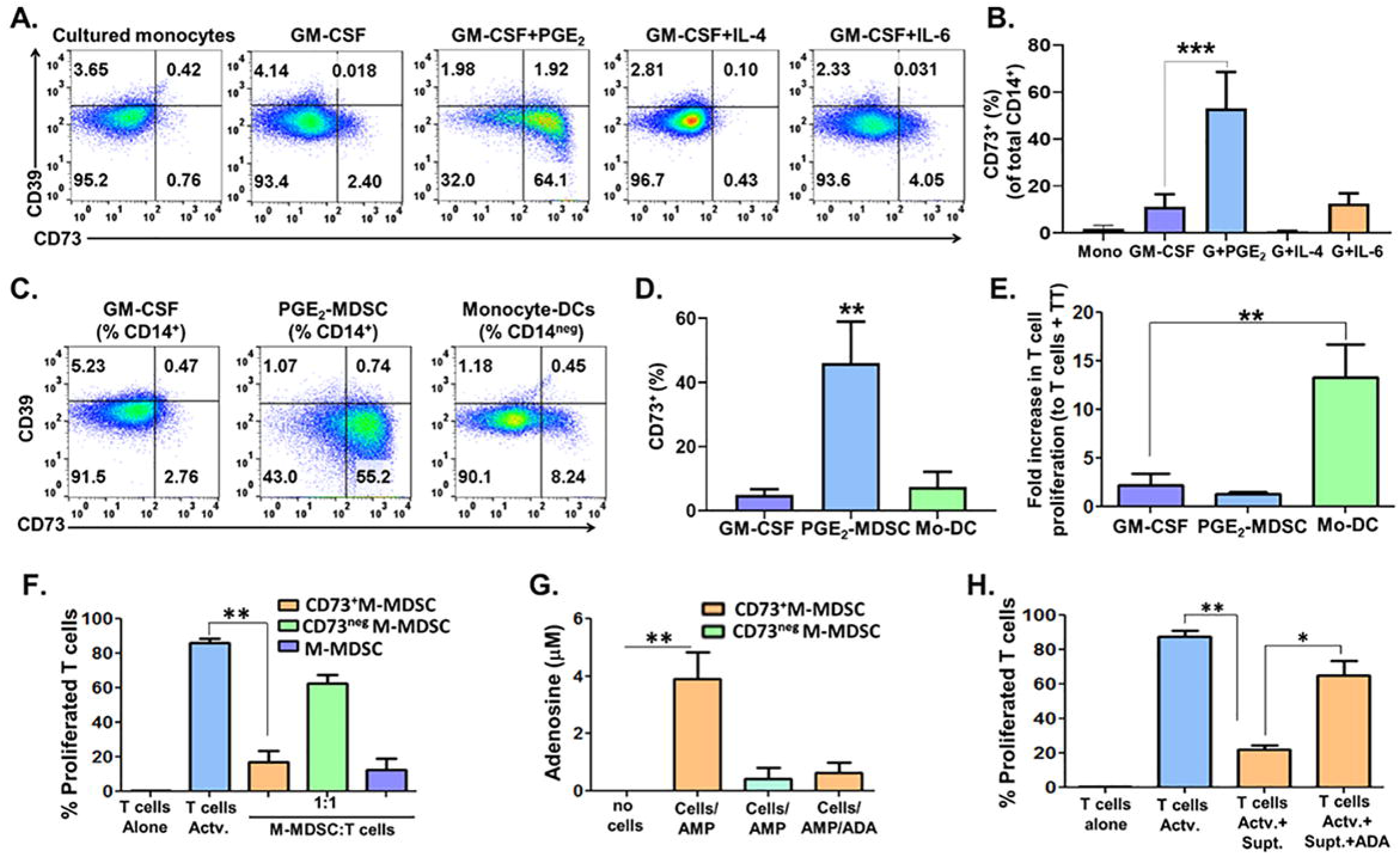
PGE_2_-induced CD73^+^ monocytic MDSCs mediate immunosuppression *via* adenosine. **A, B**. CD14^+^ monocytes from healthy donors were either untreated (cultured monocytes) or treated for 48 hours with either GM-CSF (10 ng/mL), GM-CSF and PGE_2_ (1 μM), IL-4 (10 ng/mL), or IL-6 (10 ng/mL) and CD73 expression was measured by flow cytometry. Representative dot plots **(A)** and bar graph **(B)** showing the CD73 expression (%) in the treated cells. Data, mean ± SEM of three independent experiments with monocytes from 3 different donors, *P* values: ***, p<0.0001. **C, D**. CD14^+^ monocytes from healthy donors were treated for 6 days with either GM-CSF, GM-CSF and PGE_2_ (PGE_2_-induced M-MDSCs), or GM-CSF and IL-4 (monocyte-derived dendritic cells). Dot plots **(C)** and bar graph **(D)** showing the CD73 expression (%) in GM-CSF-treated monocytes, or PGE_2_-induced M-MDSCs or in monocyte-derived dendritic cells (DCs). Data, mean ± SEM of three independent experiments with monocytes from 3 different donors, *P* values: **, p<0.001. **E**. GM-CSF-treated monocytes or PGE_2_-induced M-MDSCs or monocyte-derived DCs are analyzed for inducing autologous tetanus toxoid-specific T cell proliferation. Indicated cells were added to autologous T cells in the presence of Tetanus Toxoid (TT; 1.0 μg/mL) for 5 days and T cells proliferation was measured by incorporation of ^[3H]^thymidine. Data, mean ± SEM. *P* values: **, p<0.001. **F**. Immune suppressive activity (against anti-CD3/anti-CD28 bead activated T cells) of CD73^+^ or CD73^neg^ PGE_2_-M-MDSCs or tumor cell line-induced M-MDSCs. Data, mean ± SEM of three independent experiments, *P* values: **, p<0.001. **G**. Exogenous AMP (10 μM) or AMP and ADA (10 Units/mL) were added to fresh medium with CD73^+^ or CD73^neg^ M-MDSCs cells for 6 hours. Extracellular adenosine was measured in the culture supernatants. Data, mean ± SEM of two independent experiments, *P* values: **, p<0.001. **H**. Untreated or ADA-treated culture supernatants from CD73^+^ M-MDSCs cultured with AMP were added to T cells and suppressive activity was measured by incorporation of ^[3H]^thymidine. Data, mean ± SEM of three independent experiments, *P* values: **, p<0.001; *, p<0.05.

The addition of GM-CSF + PGE_2_ to human CD14^+^ monocytes isolated from healthy donors for a 6-day period promoted the development of CD14^+^CD33^+^HLA-DR^low/neg^ M-MDSC-like cells. A large proportion of these cells expressed CD73 (**Figs. 2C and D**) and displayed significant inhibitory activity against tetanus toxoid activated CD4^+^ T lymphocytes (**Fig. 2E**). Addition of GM-CSF alone did not induce M-MDSC differentiation (**Figs. 2C and D**). In contrast, addition of GM-CSF + IL-4 combination to monocytes for 6 days induced differentiation of monocyte-derived dendritic cells (mo-DCs) which lack CD73 expression and, expectedly, were functionally immunostimulatory and induced a significant increase in tetanus toxoid-specific T-cell proliferation (**Fig. 2E**). We further explored the immunosuppressive properties of CD73-expressing PGE_2_-M-MDSCs by determining the immunosuppressive activity of purified CD73^+^ and CD73^neg^ PGE_2_-induced M-MDSC subsets against anti-CD3/anti-CD28 antibody-activated autologous T cells. We found that the purified CD73^+^ PGE_2_-M-MDSC sub-fraction suppresses the proliferation of autologous T cells at levels comparable to that observed with tumor cell-induced M-MDSCs (**Fig. 2F**). Importantly, the CD73^neg^ PGE_2_-M-MDSC sub-fraction had substantially lower T cell inhibitory activity confirming that the CD73^+^ PGE_2_-M-MDSC sub-set mirrors the recognized suppressive activity profile of tumor M-MDSCs (**Fig. 2F**). RT-PCR analysis showed that a 4-fold increase in *A2AR* gene expression is observed in effector T cells within 48-hours of anti-CD3/anti-CD28-mediated T cell activation when compared to that in non-activated T cells (**Supplemental Fig. 2B**), corroborating the results presented in a previously published article (17). One likely scenario that rationalizes this is that CD73^+^ PGE_2_-M-MDSCs suppresses the T cell proliferation *via* the adenosine → A2AR pathway.

Next, we tested if CD73 in PGE_2_-M-MDSCs remain enzymatically active and can mediate adenosine production. CD73^+^ and CD73^neg^ PGE_2_-M-MDSCs sub-fractions were cultured in the presence and absence of AMP and adenosine levels in the supernatants was measured. CD73^+^ PGE_2_-M-MDSCs rapidly catalyzed the formation of adenosine from exogenously added AMP (**Fig. 2G**). The addition of ADA enzyme to CD73^+^ M-MDSCs cultures dramatically reduced the adenosine levels in the supernatants consistent with ADA’s ability to degrade adenosine (**Fig. 2G**). In contrast, CD73^neg^ M-MDSCs failed to generate extracellular adenosine from AMP *in vitro* (**Fig. 2G**). We then examined the T cell suppressive activity of adenosine in the CD73^+^ M-MDSCs culture supernatants. Consistent with previous studies, addition of exogenous adenosine to human CD3^+^ T cells suppressed their proliferation (**Supplemental Fig. 2C**). Importantly, we observed that the adenosine in the CD73^+^ M-MDSCs cell supernatants suppressed autologous T cell proliferation. Furthermore, addition of ADA enzyme abolished the T cell suppression mediated by CD73^+^ M-MDSCs cell supernatants (**Fig. 2H**). These data confirm that PGE_2_-induced CD73^+^ M-MDSCs actively suppress T cell activity in an adenosine-dependent manner.

Next, we delineated the molecular mechanisms involved in PGE_2_-induced CD73 expression. PGE_2_ exerts its effects by relaying signals through the EP2 and EP4 cell surface receptors (25). Both receptors are expressed on PGE_2_-MDSCs (**Supplemental Fig. 3A**). To determine which of these two receptors is primarily involved in PGE_2_-mediated CD73 expression in M-MDSCs by treating CD14^+^ monocytes with either an EP2 or EP4 agonist in the presence of GM-CSF. The EP4 agonist only modestly increased CD73 levels whereas the EP2 agonist induced CD73 expression to levels comparable to that induced by PGE_2_ (**Figs. 3A and B**). Addition of IL-4 to the EP2 agonist-treated cells completely abolished CD73 expression (**Figs. 3A and B**), suggesting that the EP2 receptor is preferentially responsible for PGE_2_-mediated CD73 induction in MDSCs. One of the effectors of EP2→PGE_2_ signaling is NF-κB (26). PGE_2_ resulted in a dramatic activation of NF-κB signaling as evidenced by an increase in phospho-NF-κB and degradation of IκBα (an inhibitor of NF-κB; **Fig. 3C**) and, as expected, resulted in an increase in CD73 expression (**Fig. 3D**) along with a sharp increase in *IL-6* expression, a downstream target of NF-κB (27, 28) (**Fig. 3D**). Addition of Forskolin, an activator of adenylate cyclase (29, 30), to the monocytes mimicked the effects of exogenous PGE_2_ on NF-κB, CD73 and IL-6 (**Figs. 3C and D**). Similarly, addition of D-cAMP, a synthetic analog of cAMP (31) induced CD73 and IL-6 to similar levels as seen with PGE_2_ (**Fig. 3E**). Taken together, these data suggest that PGE_2_ induces CD73 expression by modulating the levels of cAMP and subsequent activation of NF-κB. PGE_2_ also increased IL-10 expression which was also mimicked by Forskolin and D-cAMP (**Supplemental Fig. 3B and C**). However, while an anti-IL-10 antibody was effective at reducing PGE_2_-induced IL-10 expression, it had no effect on PGE_2_-induced CD73 expression (**Supplemental Figs. 3D and E**), suggesting that IL-10 is not involved in PGE_2_-mediated CD73 expression in M-MDSCs. While addition of IL-6 alone is not sufficient to induce CD73 expression in MDSCs (**Figs. 2A and B**), it may still be playing a role in PGE_2_-mediated CD73 induction as the IL-6 receptor (IL-6R) is expressed on both PGE_2_- and IL-6-stimulated monocytes (**Supplemental Fig. 3E**). One of the key downstream mediators of IL-6 signaling is Stat3 (32). Addition of PGE_2_ resulted in a significant activation of Stat3 which was comparable to that observed by addition of IL-6 alone (**Fig. 3F**). Inhibition of Stat3 with a Stat3 inhibitor resulted in an approximately 50% reduction in CD73 expression (**Figs. 3G, I, and J**), suggesting that PGE_2_-induced CD73 induction is, in part, mediated by Stat3 activation. Since inhibition of Stat3 only partially blocked PGE_2_-induced CD73 expression, this suggested the involvement of an additional signaling pathway(s). cAMP also activates protein kinase A which in turn activates cAMP Response Element-Binding Protein (CREB), a transcription factor (33). Addition of GM-CSF and PGE_2_ to monocytes resulted in activation of CREB (**Fig. 3H**). Inhibition of CREB with a CREB inhibitor, again reduced PGE_2_-induced CD73 by only approximately 50% (**Figs. 3H, I and J**). Addition of both the Stat3 inhibitor and the CREB inhibitor resulted in complete loss of PGE_2_-induced CD73 expression (**Figs. 3 I and J**). These results suggest that both Stat3 and CREB are required for PGE_2_-induced CD73 expression.

**Figure 3.**
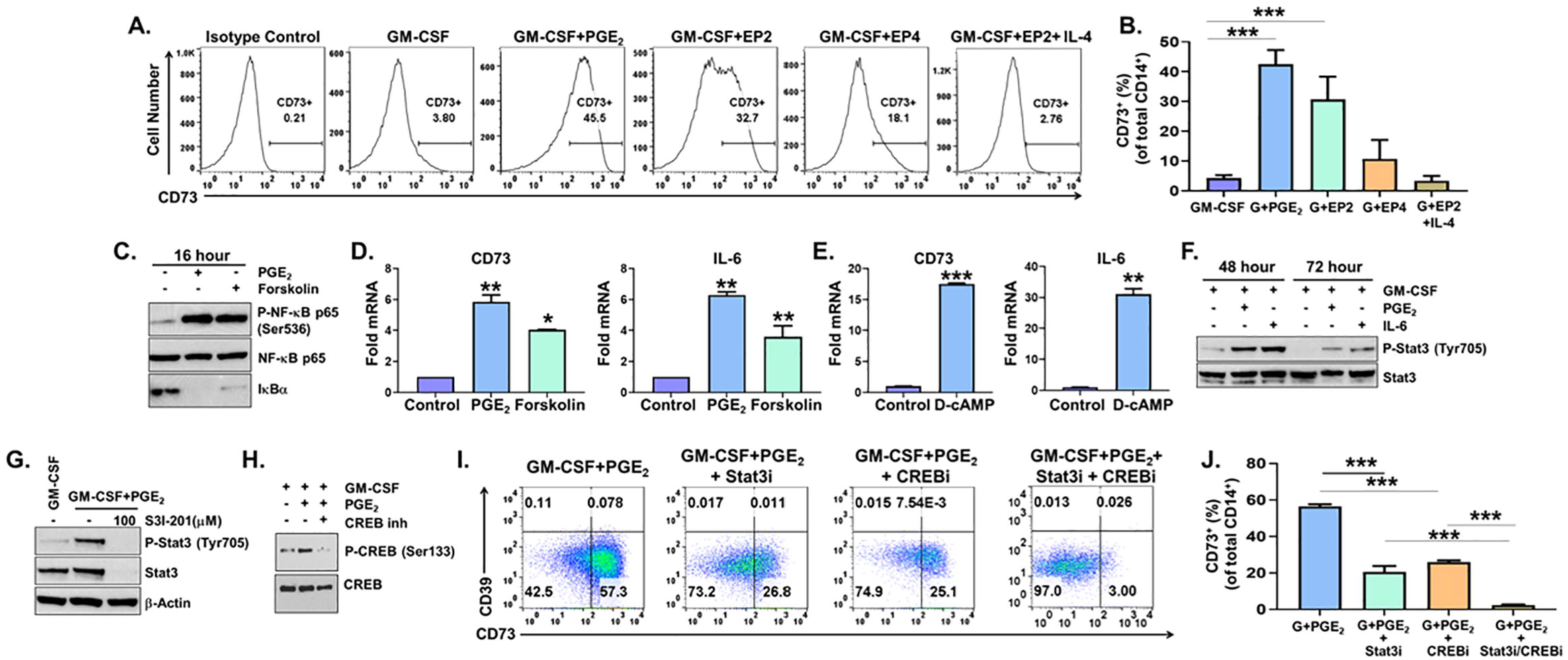
Tumor-derived PGE_2_ regulates CD73 expression in M-MDSCs *via* both a CREB and STAT3-dependent mechanism. **A, B**. EP2 receptor agonist phenocopies PGE_2_-induced CD73 expression in M-MDSCs. CD14^+^ monocytes from healthy donors were treated with either GM-CSF (10 ng/mL), GMCSF + PGE_2_ (1 μM), GM-CSF + EP2 receptor agonist (butaprost; 10 μM), GM-CSF + EP4 receptor agonist (CAY10598; 10 nM), or GM-CSF + IL-4 (10 ng/mL) + EP2 receptor agonist for 6 days, and representative histograms **(A)** and bar graph **(B)** showing the expression of CD73 in the treated cells. Data, mean ± SEM of three independent experiments, *P* values: ***, p<0.0001. **C–J**. PGE_2_ induces cell surface CD73 expression in human monocytes from healthy donors *via* a novel PGE_2_→cAMP→CREB/STAT3 pathway. **C**. CD14^+^ monocytes are treated with either vehicle (Control), PGE_2_ (1 μM), or Forskolin (10 μM) for 16 hours, and NF-κB activation measured by western blot. **D**. Bar graphs showing the CD73 mRNA and IL-6 mRNA expression in CD14^+^ monocytes treated as in C. Data, mean ± SEM, *P* values: **, p<0.001; *, p<0.01. **E**. Bar graphs showing the CD73 mRNA and IL-6 mRNA expression in CD14^+^ monocytes treated with either vehicle (Control), or D-cAMP (100 μM) for 16 hours. Data, mean ± SEM, *P* values: ***, p<0.0001; **, p<0.001. **F**. CD14^+^ monocytes were treated with either GM-CSF (10 ng/mL), GM-CSF + PGE_2_ (1 μM), or GM-CSF + IL-6 (10 ng/mL) for 48 and 72 hours, and activation of Stat3 was assessed by western blot. **G**. CD14^+^ monocytes were treated with either GM-CSF alone, GM-CSF + PGE_2_/vehicle, or GM-CSF + PGE_2_/Stat3 inhibitor (S3I-201; 100 μM) for 48 hours, and Stat3 activation was assessed by western blot. **H**. CD14^+^ monocytes are treated with either GM-CSF alone, GM-CSF + PGE_2_/vehicle, or GM-CSF + PGE_2_/CREB inhibitor (666-15; 2 μM) for 48 hours, and CREB activation was assessed by western blot. **I, J**. CD14^+^ monocytes were treated as in G and H, respectively and representative dot plots **(I)** and bar graphs **(J)** show the expression of CD39/CD73 in the treated monocytes. Data, mean ± SEM of three independent experiments, *P* values: ***, p<0.0001; **, p<0.001.

We next used murine bone marrow-derived MDSCs to test our hypothesis that tumor-derived PGE_2_ regulates T cells suppressive activity of M-MDSCs in a CD73-dependent manner. We cultured whole bone marrow (BM) cells for 5 days with GM-CSF alone, GM-CSF + PGE_2_ or GM-CSF + IL-4 and monitored the induction of CD73 in these cells. Short-term BM cultures in the presence of GM-CSF alone or GM-CSF + PGE_2_ combination resulted in the generation of cells with CD11b^hi^Gr-1^int^ M-MDSC subset as shown previously (10, 34) (**Fig. 4A**). Our data now show that both the proportions of CD73^+^ and relative expression (MFI) of CD73 was negligible in CD11b^hi^Gr-1^+^ freshly isolated BM cells (**Figs. 4A–D**). These cells did not possess T cell inhibitory activity when added at different concentrations to ovalbumin (OVA)-stimulated splenocytes obtained from OT-1 mice (**Fig. 4E**). Addition of GM-CSF to BM cells induced a moderate increase in the proportions and expression of CD73 in CD11b^hi^Gr-1^int^ cells and accordingly, displayed moderate ability to suppress the OVA antigen-specific T cell proliferation (**Figs. 4A–E**). Addition of the cytokine combination GM-CSF + IL-4 to BM cells for 5 days resulted in enrichment of CD11b^+^Gr-1^neg^CD11c^+^ dendritic cells that lacked CD73 expression and were unable to suppress OVA-antigen stimulated T cells (**Supplemental Figs. 4A–B**). In alignment with our *ex vivo* human data, treatment of BM cells with GM-CSF + PGE_2_ combination induced a significant increase in the proportions of CD73-expressing BM-MDSCs cells. These PGE_2_-induced BM-MDSCs possess superior T cell inhibitory activity (**Figs. 4A–E**). We observed that, in contrast to human monocytes, BM cells treated with GM-CSF showed increased expression of CD39 on their cell surface. However, these expression levels were not further altered with addition of PGE_2_ (**Figs. 4F and G**). To directly determine whether CD73 is functionally active, AMP was added to PGE_2_-BM-MDSCs exogenously in the presence or absence of the CD73 inhibitor AMP-CP and the culture supernatants were collected. The supernatants from untreated CD73^+^-BM-MDSCs suppressed the anti-CD3/anti-CD28-induced T cell proliferation, and CD73 inhibitor rescued the T cell proliferation (**Fig. 4H**).

**Figure 4.**
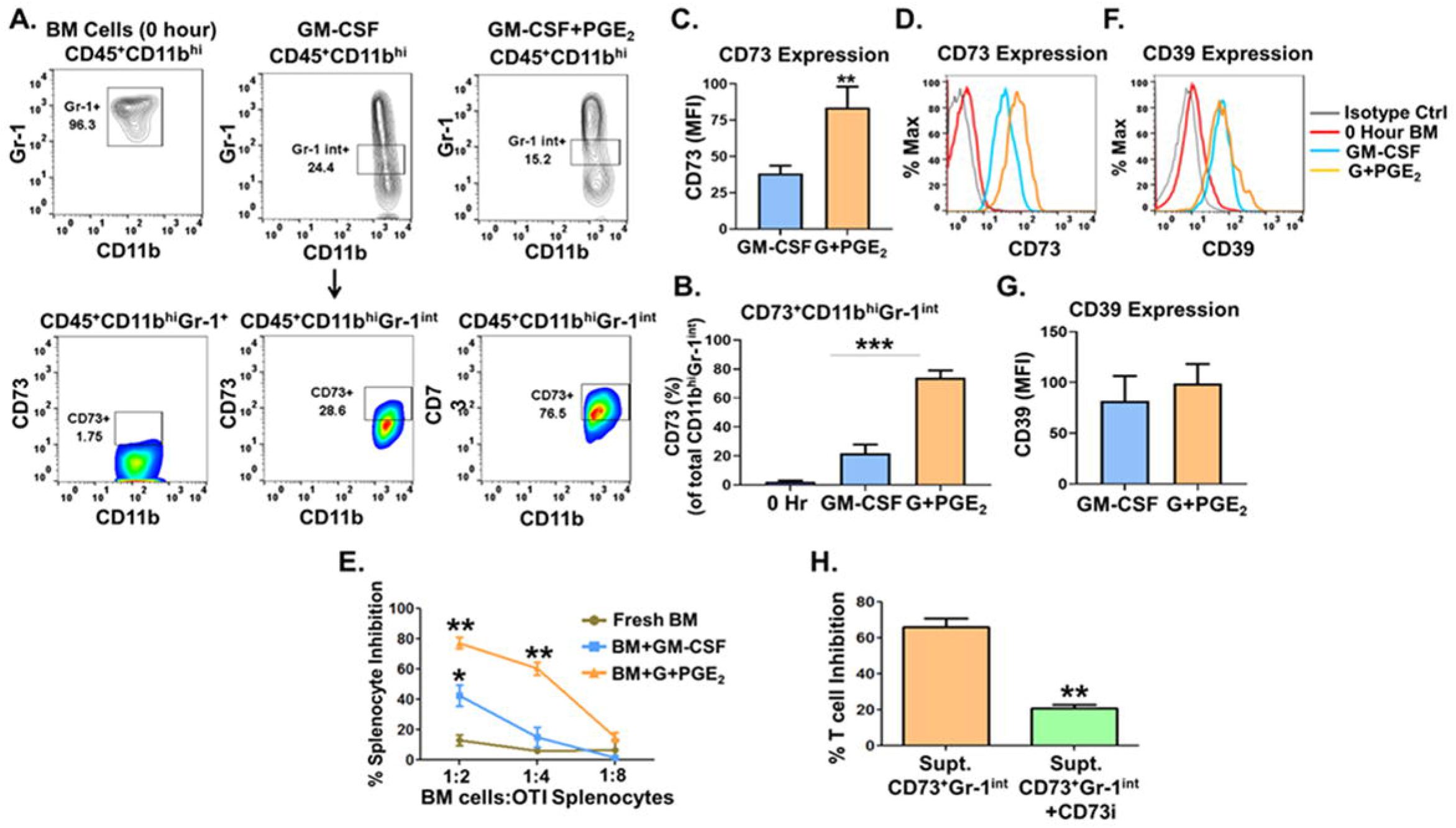
PGE_2_-induced murine bone marrow-derived CD73^+^ M-MDSCs exhibit heightened immunosuppressive activity. Murine bone marrow (BM) cells from naïve C57BL/6 were cultured with either GM-CSF (40 ng/mL) and GM-CSF and PGE_2_ (2.5 μM) for 5 days. **A–H**. Fresh BM cells or BM cells cultured with either GM-CSF or GM-CSF + PGE_2_ for 5 days were analyzed by flow cytometry for CD11b, Gr-1, CD39 and CD73 expression and T cell immunosuppressive activity. **A, B**. Accumulation of CD45^+^CD11b^hi^Gr-1^int^ M-MDSC cells and proportions of CD73^+^ cells within the CD45^+^CD11b^hi^Gr-1^int^ M-MDSC population are shown in representative contour and dot plots, respectively. **B**. Bar graphs showing proportions of CD73^+^ cells within the CD11b^hi^Gr-1^int^ population in the BM cells. Data, mean ± SEM of three independent experiments, *P* values: ***, p<0.0001. **C, D, F, and G**. Histograms and bar graphs showing the expression of CD73 **(C, D)** and CD39 **(F, G)** in total CD11b^hi^Gr-1^int^ BM cells. **E**. Fresh BM cells or BM cells were cultured with GM-CSF or GM-CSF + PGE_2_ for 5 days. Cells were harvested and added to splenocytes purified from OT-I transgenic mice at 1:2, 1:4 and 1:8 ratio in the presence of ovalbumin (250 μg/mL) per well for 4 days. T cell proliferation was measured by incorporation of ^[3H]^thymidine. Data, mean ± SEM of three independent experiments, *P* values: **, p<0.001; *, p<0.01. **H**. The effect of CD73 inhibitor AMP-CP on the *in vitro* suppressive activity of CD73^+^ BM-derived MDSCs. CD73^+^ BM-MDSCs were purified and cultured with AMP in the presence or absence of CD73 inhibitor AMP-CP (10 μM) and supernatants were collected and added to T cells in the presence of anti-CD3 and anti-CD28 antibodies, and T cell proliferation was measured by incorporation of ^[3H]^thymidine. Data, mean ± SEM of three independent experiments, *P* values: **, p<0.001.

To validate our *ex vivo* data in a localized tumor microenvironment *in vivo* in mice, we initially measured PGE_2_ levels *in vitro* in a panel of three syngeneic immunocompetent mouse tumor models— B16 melanoma, Lewis lung carcinoma (LLC), and 4T1 breast carcinoma. Using ELISA, we measured the baseline levels of PGE_2_ produced by the tumor cell lines. The data showed that B16-F10 melanoma and LLC cells produce low levels of PGE_2_, while 4T1 cells produce high levels of PGE_2_ into the culture supernatants (**Fig. 5A**). To confirm our hypothesis that tumor-derived PGE_2_ influences CD73 expression in myeloid cells, we stably knocked down *PTGES* (prostaglandin E synthase, an enzyme critical for the synthesis of PGE_2_) in 4T1 cells (4T1-*shPTGES*) using shRNA technology, thereby, blocking PGE_2_ production in the tumor cells (**Fig. 5A**). We then established subcutaneous B16-F10, LLC, 4T1-shScr and 4T1-*shPTGES* tumors in syngeneic mice. Tumors were harvested 18–21 days post-tumor cell injection and the CD73 expression was profiled in MDSCs present in the tumor digests. Our data show that the frequencies of CD73-expressing CD11b^hi^GR-1^int^ M-MDSCs are significantly higher in tumors with higher PGE_2_ levels (**Figs. 5B-D**); we observed a strong correlation between PGE_2_ levels and CD73^+^ M-MDSC frequencies in these tumor models (**Fig. 5E**). We also compared the CD73 expression levels between CD11b^hi^Gr-1^int^ M-MDSCs and CD11b^hi^Gr-1^hi^ G-MDSCs (10) subsets in 4T1 tumors. The G-MDSC sub-population expressed low levels of CD73 (**Fig. 5F**) and as such, no correlation was observed between PGE_2_ levels and CD73 expressing G-MDSCs present in the tumors. These data indicated that the induction of CD73 expression on M-MDSCs but not on PMN-MDSC was reliant on PGE_2_ expression from within the tumors.

**Figure 5.**
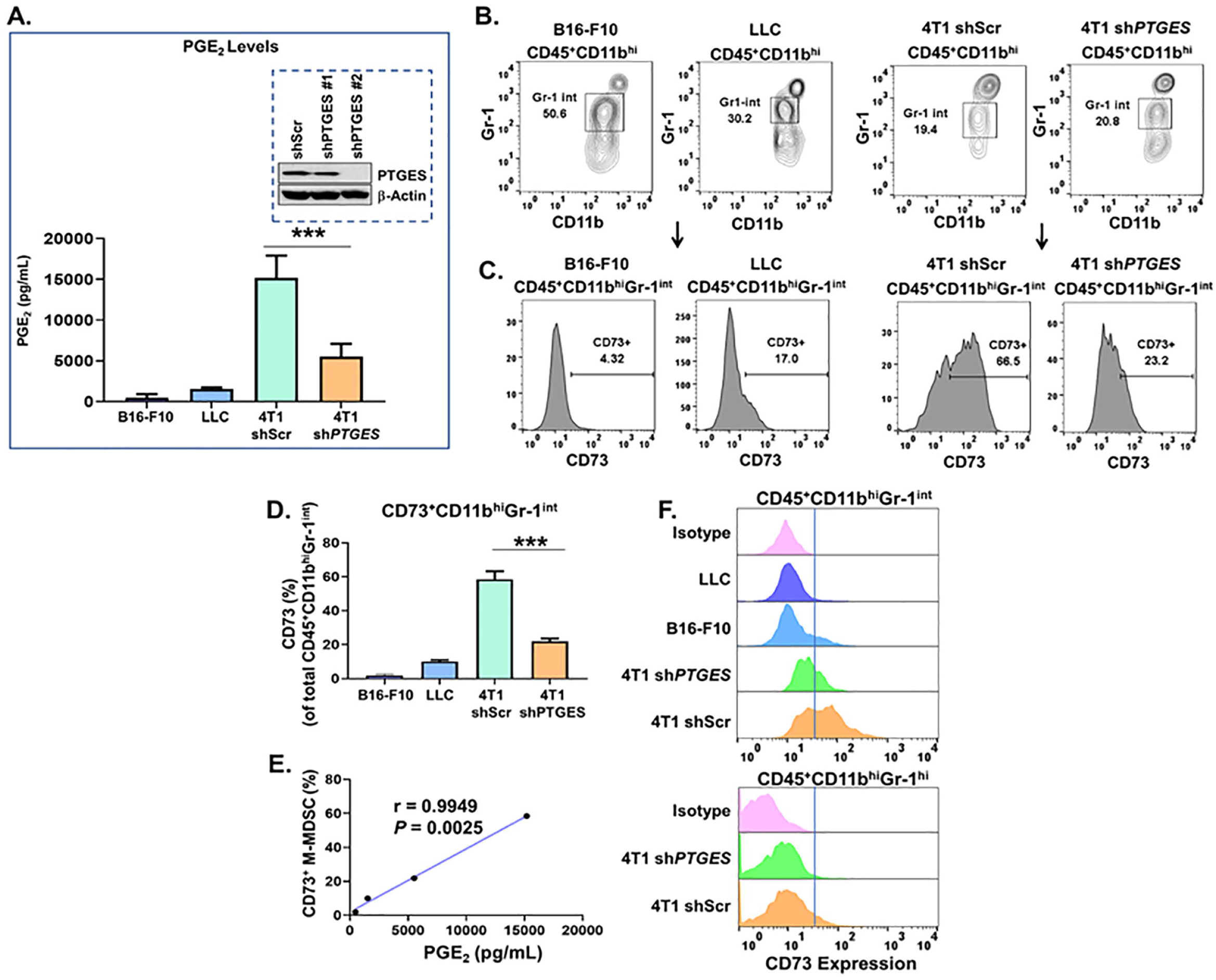
Genetic depletion of tumor-derived PGE_2_ reduces CD73 expression on monocytic MDSCs in the tumor microenvironment. **A**. Bar graphs showing PGE_2_ levels culture supernatants of murine B16-F10 melanoma cells, LLC lung cancer cells, and 4T1 breast cancer cells with either *PTGES* knockdown or scrambled control (4T1*shPTGES*, 4T1shScr, inset) as measured by ELISA (n = 2 independent ELISA expt.). Data, mean ± SEM, *P* values: ***, p<0.0001. **B–E**. Syngeneic tumors implanted subcutaneously into BALB/c mice (4T1*shPTGES* and 4T1shScr cells) and C57BL/6 mice (B16-F10 and LLC cells) (n = 5 mice per tumor type). Tumors were isolated (200–300mm^3^), digested and stained for CD45, CD11b, Gr-1, and CD73 expression. **B-D**. Dot plots showing the proportions of CD45^+^CD11b^hi^Gr-1^int^ M-MDSCs **(B)**, and histograms **(C)** and bar graphs **(D)** showing % CD73^+^ cells within the CD45^+^CD11b^hi^Gr-1^int^ M-MDSC population in the implanted tumors. Data, mean ± SEM, *P* values: **, p<0.001. **E**. Correlation between intra-tumoral % CD73^+^ M-MDSCs *vs* PGE_2_ levels produced by the corresponding tumor cells. *p* = 0.0050, r^2^ = 0.9950. **F**. Histograms showing CD73 expression in the CD45^+^CD11b^hi^Gr-1^int^ M-MDSC and CD45^+^CD11b^hi^Gr-1^hi^ G-MDSC population in the implanted tumors.

To evaluate the *in vivo* functional contributions of the PGE_2_ synthesis pathway to the expansion of immunosuppressive CD73^+^ M-MDSCs in a physiologically relevant environment, we established an orthotopic KP1.9 lung adenocarcinoma mouse model. KP1.9 cells are originally isolated from *LSL-Kras*^*G12D*^: *p53*^*fl/fl*^ (KP) transgenic mouse model of non-small cell lung cancer (NSCLC) (35). Our data from the orthotopic KP1.9 mouse model show that tumor development in the mice lungs initiates around day 24 post-tumor cell injection and progressively intensified until day 40–42; (**Figs. 6A and B**). PGE_2_ levels were high in these tumors with an average of 189 ± 48 ng of PGE_2_ present per gram of tumor tissue (**Fig. 6C**). We also found that a significant percentage of M-MDSCs expressed CD73 in the KP tumor-bearing lungs when compared to a similar population of monocytes present in lungs obtained from naïve mice (**Figs. 6D, E**). CD73-M-MDSC functionality in KP lung tumors was determined using the OT-I TCR transgenic mice and OVA antigen presentation. Our data indicate that CD73-expressing KP lung tumor-derived M-MDSCs significantly suppressed OVA-antigen specific CD8^+^ T cell proliferation, while monocytes derived from naïve lungs failed to exhibit any T cell suppressive activity (**Fig. 6F**). In addition, AMP-containing culture supernatants from the CD73^+^ M-MDSCs from KP mice mediated potent suppression of anti-CD3/anti-CD28-induced T cell proliferation (**Fig. 6G**). We validated our observations in CD73-deficient mice (Nt5e^−/−^). Orthotopic KP lung tumors were established in wild-type and Nt5e^−/−^ mice (**Fig. 6H**) and found that M-MDSCs from Nt5e^−/−^ were less immunosuppressive than wild-type M-MDSC cells (**Fig. 6I, J**).

**Figure 6.**
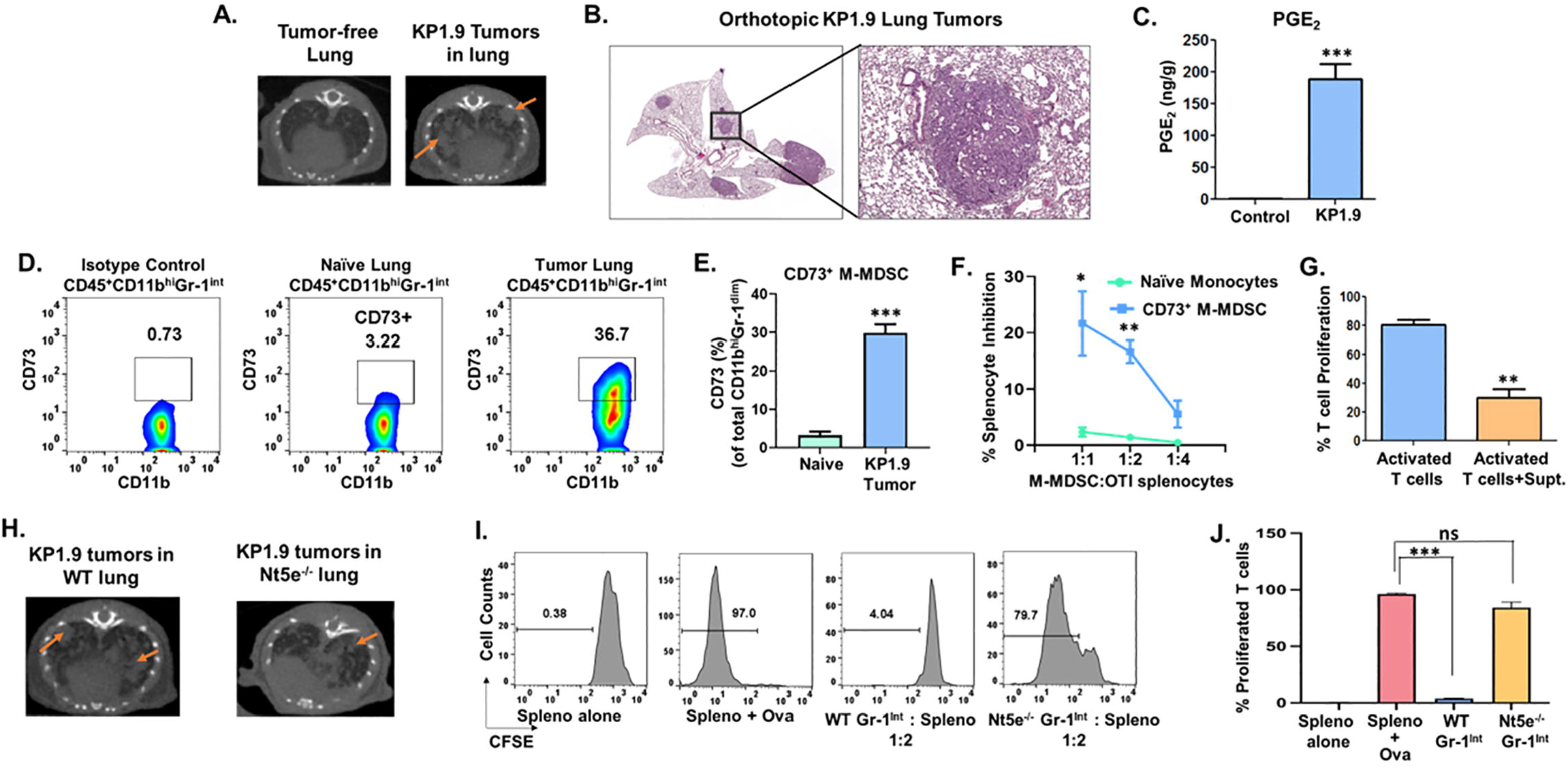
Genetic deletion of CD73 reduces the adenosine-dependent suppressive activity of monocytic MDSC in the tumor microenvironment. **A–G**. C57BL/6 mice (n = 20 mice) were injected with KP1.9 lung adenocarcinoma cells (5 × 10^5^) *via* intravenous (i.v.) route. **A**. Representative micro-CT images of lungs of representative KP1.9 mice, imaged at day 40. Red arrowheads point to an individual tumor. **B**. Mice were sacrificed after 40 days and lungs were perfused and fixed. Hematoxylin and eosin staining (H&E) stained and digitally-scanned section of KP1.9 tumor-bearing mouse lung. Magnification: 1× and 8×. **C**. Murine KP1.9 tumor-bearing lungs were harvested and PGE_2_ levels in the lung tissue were measured using mass spectrometry. Bar graph shows the PGE_2_ levels in KP1.9 tumor-bearing lung tissue (n = 4 mice). Data, mean ± SEM, *P* values: ***, p<0.0001. **D–F**. Lungs from naïve and KP1.9 tumor-bearing mice (n = 9 mice per group from 3 independent experiments) were processed into single-cell suspensions and immune cells were isolated using percoll gradient. Dot plots **(D)** and bar graphs **(E)** showing % CD73^+^ cells within the CD45^+^CD11b^hi^Gr-1^int^ population in the naïve and KP1.9 tumor-bearing mice lungs. **F**. CD73^+^ CD45^+^CD11b^hi^Gr-1^int^ cells from lung tissue obtained from tumor-bearing and naïve mice (from n = 6 mice per group) were purified and co-cultured with OT-I splenocytes and ovalbumin (250 μg/mL) for 4 days. T cell proliferation was measured using ^[3H]^-thymidine. Data, mean ± SEM, *P* values: ***, p<0.0001; **, p<0.001; *, p<0.01. **G**. Exogenous AMP (10 μM) were added to fresh medium with CD73^+^ M-MDSCs cells for 6 hours. Culture supernatants from CD73^+^ M-MDSCs/AMP cultures were added to T cells and proliferation was measured by incorporation of ^[3H]^thymidine. Data, mean ± SEM of two independent experiments, *P* values: **, p<0.001. **H–J**. Wild-type (C57BL/6) and Nt5e^-/-^ mice (n = 10 mice/group) were injected with KP1.9 lung adenocarcinoma cells (5 × 10^5^) *via* intravenous (i.v.) route. **H**.Representative micro-CT images of lungs of representative KP1.9 mice, imaged at day 40. Red arrowheads, individual tumor. **I, J**. Lungs from WT and Nt5e^-/-^ KP1.9 tumor-bearing mice were processed into single-cell suspensions and CD11b^hi^Gr-1^int^ M-MDSCs were purified. M-MDSCs were co-cultured with CFSE-labeled splenocytes from OT1 transgenic mice and ovalbumin (250 μg/mL) for 4 days. CD8^+^ T proliferation was measured using flow cytometry. Histograms **(I)** and bar graph **(J)** showing the % proliferation of T cells. Data, mean ± SEM of two independent experiments (n = 6 mice), *P* values: ***, p<0.001.

We propose a novel immunotherapeutic strategy involving administration of adenosine deaminase (ADA)—an enzyme that specifically and irreversibly converts T cell suppressive adenosine into the non-immunosuppressive nucleoside inosine. The PEGylated version of ADA (PEG-ADA) is an FDA-approved biologic that is currently used as an enzyme replacement therapy for ADA deficiency in pediatric SCID patients (20). PEG-ADA is a conjugate of numerous strands of monomethoxy PEG (5000 Da) covalently attached to the enzyme ADA. The enzyme ADA is involved in purine metabolism; it breaks down adenosine from food and is required for the turnover of nucleic acids in tissues (36, 37). We have begun to test the idea that enzymatic conversion of adenosine to inosine in tumors might facilitate their destruction by CD8^+^ T cells. To evaluate whether enzymatic conversion of adenosine to inosine is a viable immunotherapeutic approach, we initially assessed the potential efficacy of PEG-ADA to induce immune cell-mediated rejection of lung tumors using an implantable KP1.9 lung adenocarcinoma mouse model (35). C57BL/6 mice were injected subcutaneously with KP1.9 cells and PEG-ADA or PBS (vehicle control) was injected intraperitoneally (*i*.*p*.) on day 0 and then every two days for 28 days. As shown in **Fig. 7A**, KP1.9 tumors in PEG-ADA-treated mice were significantly smaller and grew at a much slower rate than those in control mice. Because extracellular adenosine promotes evasion from anti-tumor T cell responses, we set out to determine whether PEG-ADA-mediated catabolism of intra-tumoral adenosine might alter pro-tumorigenic T cell responses in tumor bearing mice. We assessed the quantity and quality of anti-tumor immune cell responses generated within the KP1.9 tumors in the PEG-ADA-treated and vehicle-treated control mice. For analysis of tumor-infiltrating immune cells, tumors from drug-treated and control mice were excised at the end of therapy and analyzed for expression of surface and functional markers of T cells by FACS. Our data indicate that significantly higher frequencies of IFN-γ producing CD8^+^ T cells were obtained from PEG-ADA-treated tumors than that observed in the vehicle-treated tumors (**Fig. 7B**) while intra-tumoral adenosine levels were reduced (**Fig. 7C)**. These results suggest that PEG-ADA treatment reduces intratumoral adenosine levels and, consequently, generates anti-tumor T cell responses. This illustrates the important point that PEG-ADA may also be useful to treat tumors with high baseline levels of PGE_2_/adenosine. Importantly, PEG-ADA functionally synergizes with anti-PD-1 Ab to reduce KP1.9 cells’ resistance to anti-anti-PD-1 monotherapy resulting in significantly reduced subcutaneous tumor burden (**Fig. 7D**) and increased survival (**Fig. 7E**). suggesting that PEG-ADA therapy enhances anti-PD-1 antibody therapeutic efficacy.

**Figure 7.**
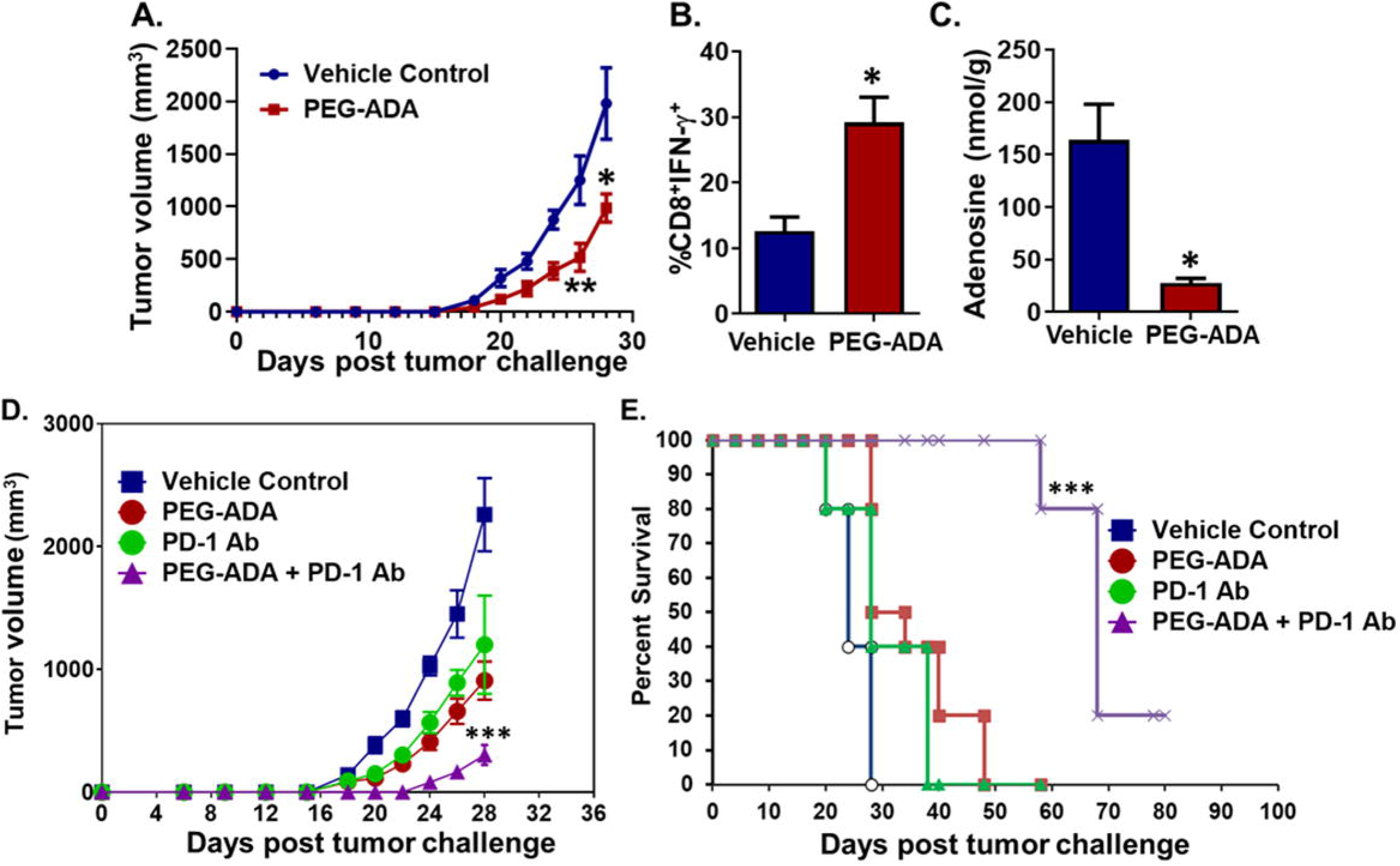
PEG-ADA depletes intra-tumoral adenosine and enhances tumor response to anti-PD-1 immunotherapy. **A**. C57BL/6 mice were injected subcutaneously (s.c.) with KP1.9 lung adenocarcinoma cells (2 × 10^5^ cells). PEG-ADA (2 Units/mouse) or PBS (vehicle) was injected on day 0 and then every other day *via* intraperitoneal (i.p.) route and tumor volume was measured. Data, mean ± SD of n = 20 mice per group, *P* values: *, p≤0.05; **, p≤0.005; relative to control group. **B**. PEG-ADA treatment significantly increases percentages of IFN-γ expressing CD8^+^ T cells. Mice were euthanized 28 days after tumor challenge, tumors were removed and enzymatically digested. Tumor-infiltrating cells from PEG-ADA and vehicle-treated mice were stimulated with KP1.9 cell lysate (50 μg/mL) for 24 hours. Cells were then restimulated for 6 hours with KP lysate (50 μg/mL) in the presence of Brefeldin A (1 μL/mL of the culture medium). After restimulation, cells were stained for surface expression of CD8 and intracellular expression of cytokines and analyzed by flow cytometry. Bar graphs showing percentages of CD45^+^CD3^+^CD8^+^IFN-γ^+^ in tumors derived from vehicle- and PEG-ADA-treated mice (n = 5 mice/treatment group). Results are expressed as percentages of total CD45^+^ cells. Data, mean± SEM of n = 6 mice per group. *P* values: *, p≤0.05. **C**. PEG-ADA treatment depletes intra-tumoral adenosine levels in KP1.9 tumors. Bar graphs showing adenosine levels in KP1.9 tumors derived from vehicle- and PEG-ADA-treated mice. Data, mean ± SEM of n = 5 mice per group, *P* values: *, p≤0.05. **D, E**. C57BL6 mice were injected with KP1.9 lung adenocarcinoma cells at 2 × 10^5^ cells per mouse *via* the s.c. route. Four groups of 8 tumor-bearing mice/group received route the following treatments *via* the i.p.: 1) Vehicle control (PBS); 2) anti-PD-1 alone (clone RMP1-14; 250 μg/mouse, i.p.); 3) PEG-ADA alone (2 Units/mouse; i.p. route); and 4) anti-PD-1 plus PEG-ADA. **D**. Tumor sizes were plotted as tumor volume/time. Data, mean ± SD of n = 8 mice per group, *P* values: ***, p<0.0001; relative to vehicle-treated group. **E**. Kaplan-Meier survival curves of treated mice. ***, p<0.0001; log-rank test.

## Discussion

ICIs have significantly improved the clinical management of cancer, leading to the approval of several PD-1 or PD-L1 antibodies as therapeutic drugs both in the first- and second-line settings. Despite the improvements, immune evasion limits the efficacy of such immunotherapies. Monocytic MDSC-mediated suppressive activity is hypothesized to be one of the more prevalent mechanisms that provides an obstacle for immune-based therapeutic intervention (38). MDSCs inhibit anti-tumor T cell activity, thereby promoting an immunosuppressive milieu within the TME (12, 39). Not surprisingly, advanced cancer patients with higher levels of MDSCs prior to treatment display a lower response rate to ICI therapy (13, 40, 41). We show here that a prominent mechanism for the creation of the immunosuppressive environment within the tumors involves the PGE_2_ → CD73^+^ M-MDSC → Adenosine pathway. Our data show that the tumor cell-derived PGE_2_ regulates CD73 expression in M-MDSCs and that CD73 expression sustains the M-MDSC function leading to adenosine-dependent inhibition of CD8^+^ T cell activation. Furthermore, we illustrate a treatment strategy that repurposes a clinically tested drug that achieves regression of most of the tumors in an anti-PD-1 treatment-resistant, clinically relevant lung cancer mouse model; we demonstrate that enzymatic depletion of intra-tumoral adenosine with PEG-ADA improves the therapeutic efficacy of anti-PD-1 antibody.

PGE_2_ is a proinflammatory molecule produced by cancer and stromal cells (42). In the present study, we have identified PGE_2_ as a primary tumor-derived inflammatory mediator responsible for the up-regulation of CD73 expression in CD14^+^ myeloid cells within the TME. Overproduction of PGE_2_ is a hallmark of many human malignancies (43-45), therefore, the currently identified PGE_2_ → CD73 mechanism is likely to be present in multiple inflammation-associated cancers. We and others have shown that PGE_2_ is a critical mediator of M-MDSC expansion (9, 34, 46). A recent study has shown that oncolytic virus-mediated targeting of tumor PGE_2_ sensitizes murine breast tumors to immunotherapies by reducing G-MDSC infiltration into the TME (45). Interestingly, in our *ex vivo* and *in vivo* functional studies, tumor-derived PGE_2_ sustains M-MDSC’s suppressive activity in a CD73-dependent manner but does not affect CD73 expression on G-MDSCs. Additionally, PGE_2_ is known to redirect the differentiation of human DCs toward functionally stable MDSCs (9, 46) and as such, DCs and M-MDSCs have opposing immune functions. In line with these observations, our data indicate that GM-CSF/IL-4-driven monocyte-derived DCs lack CD73 expression and addition of IL-4 to monocyte cultures leads to repression of EP2 agonist-induced CD73 expression in these cells.

Apoptotic T and NK regulatory cells in the TME have been shown to express CD73 and inhibit T cell responses *via* the production of adenosine (47, 48), however, very little is known about CD73 expression in other immune cell populations that infiltrate the tumor microenvironment. The limited number of studies that have investigated the involvement of CD73 on MDSC-mediated immunosuppression are largely focused on the effects of this enzyme in granulocytic MDSCs (G-MDSCs) (49-51). Additionally, the molecular mechanisms involved in the regulation of the CD73 ectonucleotidase are not well understood (52). Previously published reports have focused solely on the transcription factor Stat3 (48, 53). Our data suggest a novel signaling pathway regulating CD73 expression in M-MDSCs that involves both the transcription factors Stat3 and CREB (**Fig. 8**). This new signaling pathway is initiated by tumor-derived PGE_2_ that transduces intracellular signals through the EP2 receptor on M-MDSCs and involves a potential IL-6-dependent feed forward loop activating Stat3, which together with cAMP → PKA-activated CREB, induce maximal CD73 expression. This rational conclusion is supported by the fact that the CD73 promoter is strongly predicted to contain binding sites for both Stat3 and CREB, as determined by MatInspector, a software tool that utilizes a large library of matrix descriptions for transcription factor binding sites to accurately locate sites within DNA sequences to which transcription factor can bind. (https://www.genomatix.de/cgibin/matinspector_prof/mat_fam.pl?s=bd56c95a6d3b2baa23cdd83f6acfc74b). The increase in the levels of CD73 expression on M-MDSCs results in an increased accumulation of adenosine in the TME, which ultimately suppresses anti-tumor T cell activity. We believe that there is no PGE_2_ feed forward loop in MDSCs since exogenous PGE_2_ induces IL-10 expression which downregulates COX-2 (54, 55), and COX-2 is required for PGE_2_ synthesis (56). The expression of CD73 is sustained by the exposure of the M-MDSCs to the continuous supply of tumor cell-derived PGE_2_ in the TME.

**Figure 8.**
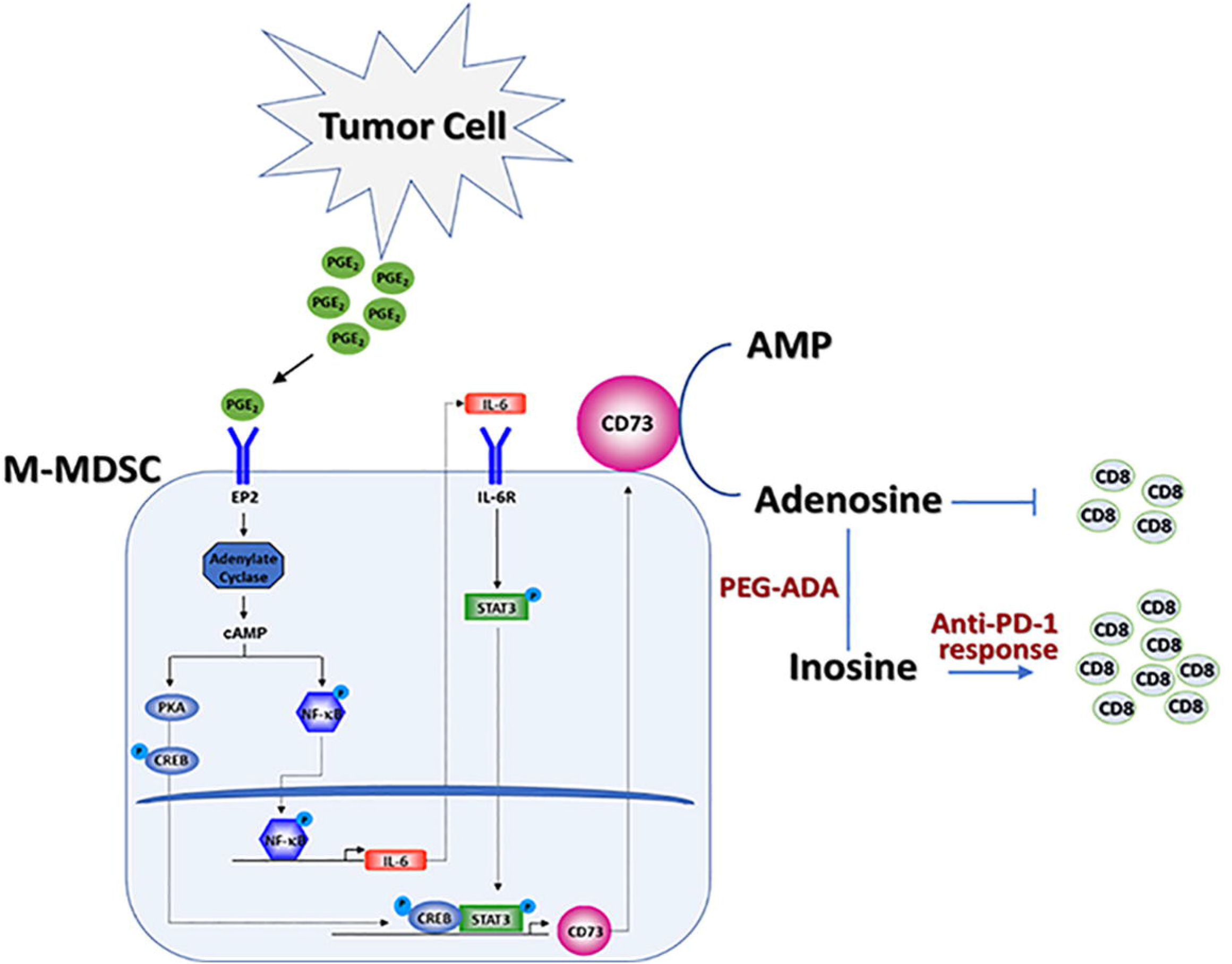
Model of the proposed mechanism of M-MDSC suppressive activity. Tumor-derived PGE_2_ initiates a signaling cascade in M-MDSCs resulting in Stat3/CREB-dependent upregulation of CD73 expression, inducing increased levels of adenosine in the TME culminating in the inhibition of anti-tumor T cell activity and thus, limits the efficacy of immunotherapy. Depletion of adenosine with PEG-ADA will overcome adenosine-mediated immunosuppression and sensitize tumors to immunotherapy.

Our data suggest that circulating CD14^+^ monocytes from NSCLC patients lack CD73 expression on their cell surface. It is possible that CD73 expression could be upregulated in these circulating monocytes upon their infiltration into the PGE_2_-enriched TME. These data also highlight the diversity in the effector functions of circulating and tumor-infiltrating myeloid cells. Intriguingly, high levels of soluble CD73 in the blood have been shown to correlate with poor prognosis in metastatic cancer patients (57). CD73 is an enzyme that hydrolyzes extracellular AMP when expressed on the cell surface, however, there is a possibility that tumor-educated M-MDSCs present in the TME can shed CD73 protein into the extracellular milieu. Alternatively, MDSCs have been shown to secrete exosomes (58) which could be loaded with CD73 molecules. These possibilities remain to be explored. Our results also indicate that human M-MDSCs express CD73 but not CD39; this is in contrast with that observed in murine M-MDSCs, which express both CD39 and CD73 ectoenzymes. These data reiterate the previously observed differences between mouse and human MDSCs (59-61), and provide additional evidence that despite the phenotypic diversity, a similar PGE_2_-induced molecular program controls the CD73-adenosine immunoregulatory axis in both human and murine M-MDSCs.

Despite the prominence of adenosine in driving cancer progression, no efficacious therapeutic strategy that specifically targets adenosine in solid tumors is currently available (62). Recently, the clinical activity of an A2AR antagonist (CPI-444) administered either alone or with anti-PD-L1 was evaluated in patients with a spectrum of advanced tumors including NSCLC; disappointingly, complete responses were not recorded in any tumor type (63). Humanized anti-CD73 antibodies are currently being tested in clinical trials. CPI-006, a humanized IgG1 FcγR binding-deficient antibody is currently being tested as a monotherapy and in combination with CPI-444 for solid tumors (NCT03454451). A Phase 1 study of the anti-CD73 antibody, IPH5301, is currently being performed in patients with advanced solid tumors (NCT05143970). Results from these trials are anticipated. We tested a novel immunotherapeutic strategy involving administration of adenosine deaminase (ADA), a highly efficacious, long lasting and non-toxic adenosine depleting enzyme. Use of ADA represents a unique and highly clinically relevant approach to disengage adenosine-mediated T cell immune suppression and overcome anti-PD-1 therapeutic resistance in cancer patients. ADA will not only eliminate the adenosine present in the TME, but also converts it to inosine, which can be used by T cells as an energy source to support their effector functions in a glucose-deprived TME (64). Importantly, inosine has been shown to enhance the efficacy of immunotherapy by activating anti-tumor T cells in a A2AR-dependent manner (65) in mouse models of cancer (64, 65). The PEGylated version of ADA (PEG-ADA) is an FDA-approved biologic that is currently used as an enzyme replacement therapy for ADA deficiency in pediatric SCID patients (20). In tumors, as in SCID patients, adenosine accumulates and inhibits immune rejection. The attachment of PEG to the ADA molecule slows the clearance of ADA, increases the circulation half-life, and lowers binding by host antibodies, thus reducing toxicity (66). When injected systemically, PEG-ADA irreversibly eliminates all available extracellular adenosine by maintaining a high level of ADA enzyme activity for a prolonged period, thereby reducing immune evasion and correcting impaired T cell function. PEG-ADA enzyme can effectively eliminate the end-product of all the CD73 nucleotidase activity, i.e. adenosine, in the TME and thus can be more efficacious in targeting both intrinsic and acquired therapeutic resistance to ICIs when compared to that offered by CD73-targeting antibodies.

There were certain limitations to our study. It is unclear whether KP tumor regression in response to the PEG-ADA/anti-PD-1 combination therapy was CD73^+^ MDSC-dependent. Additionally, we were not able to formally determine the specific contribution of CD73^+^ M-MDSCs to the total accumulated adenosine levels in the TME. Both of these limitations will be addressed in future studies using genetic deletion of CD73 specifically in M-MDSCs. Finally, our murine studies are limited to orthotopic primary tumor settings and lack information regarding metastatic disease setting which will be investigated in future studies using multiple tumor models.

In summary, PGE_2_ produced by tumor cells regulates CD73 expression in M-MDSCs *via* a novel Stat3 and CREB-dependent pathway resulting in increased extracellular adenosine generation that suppresses CD8^+^ T cell activation in solid tumors. Furthermore, enzymatic depletion of extracellular adenosine reverses CD8^+^ T cell exhaustion and sensitizes tumors to anti-PD-1 therapy.

