## Supplemental Figures for "Tumor-educated monocytes suppress T cells *via* adenosine and depletion of adenosine in the tumor microenvironment with adenosine deaminase enzyme promotes response to immunotherapy"

### Supplemental Figure 1

**A.**

| Gene | Fold Change | *P*-Value |
|---|---|---|
| | Mono vs. M-MDSC | Mono vs. M-MDSC |
| **CD73** | **16.654** | **7.23E-05** |
| **CD39** | **-3.3516** | **0.0503** |
| IL-6 | 6.32013 | 2.56E-06 |
| VEGF A | 3.44231 | 1.11E-05 |
| ARG2 | 5.70656 | 1.01E-06 |
| IDO1 | 1.88184 | 0.0004 |
| NOX4 | 7.2177 | 3.99E-08 |

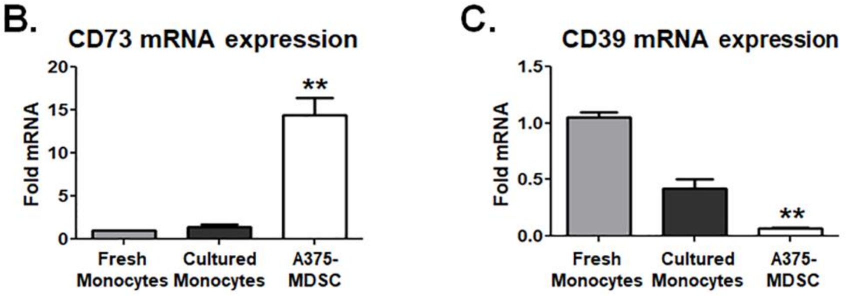


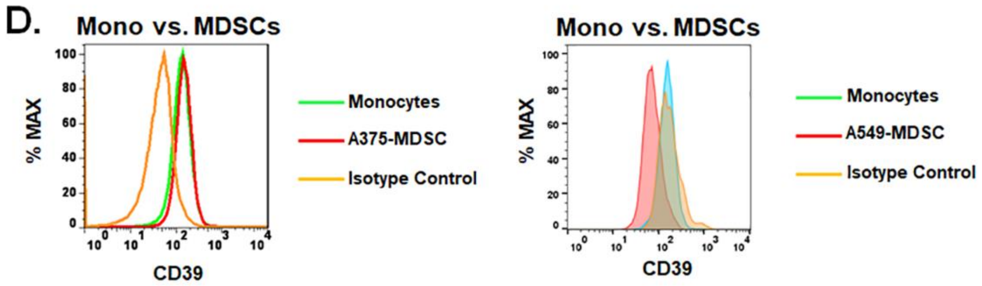


**Supplemental Figure 1. A–C.** Microarray analysis **(A)** and quantitative RT-PCR analysis show that CD73 mRNA expression is increased **(B)** and CD39 mRNA expression is decreased **(C)** when compared to the expression levels in cultures monocytes. **D.** Representative histograms showing the expression of CD39 in A549 and A375 tumor cell-induced human M-MDSCs and cultures monocytes alone. Data, mean ± SEM of three independent experiments with monocytes from 3 different donors, *P* values: **, p<0.001.

### Supplemental Figure 2

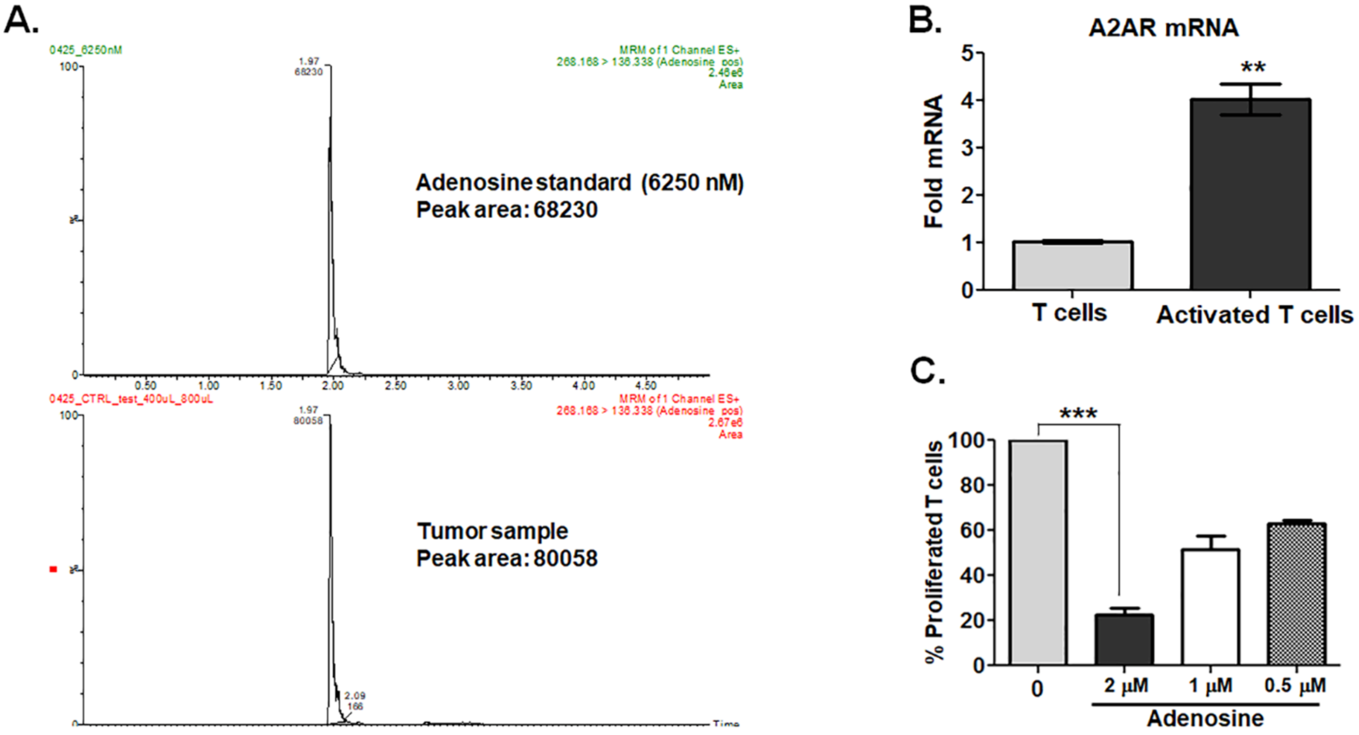


**Supplemental Figure 2. A.** Ion chromatogram showing the adenosine in reference standard and in the test tumor sample. **B.** Quantitative PCR analysis of A2AR mRNA in human T cells alone or in T cells that were activated with anti-CD3/anti-CD28 antibodies (n = 3). Data, mean ± SEM, *P* values: **, p<0.001. **C.** Human T cells were activated with anti-CD3 and anti-CD28 in the absence or presence of different doses of adenosine and proliferation was measured using [3H]-thymidine incorporation. Data, mean ± SEM, P values: ***, p<0.001.

#### Supplemental Figure 3

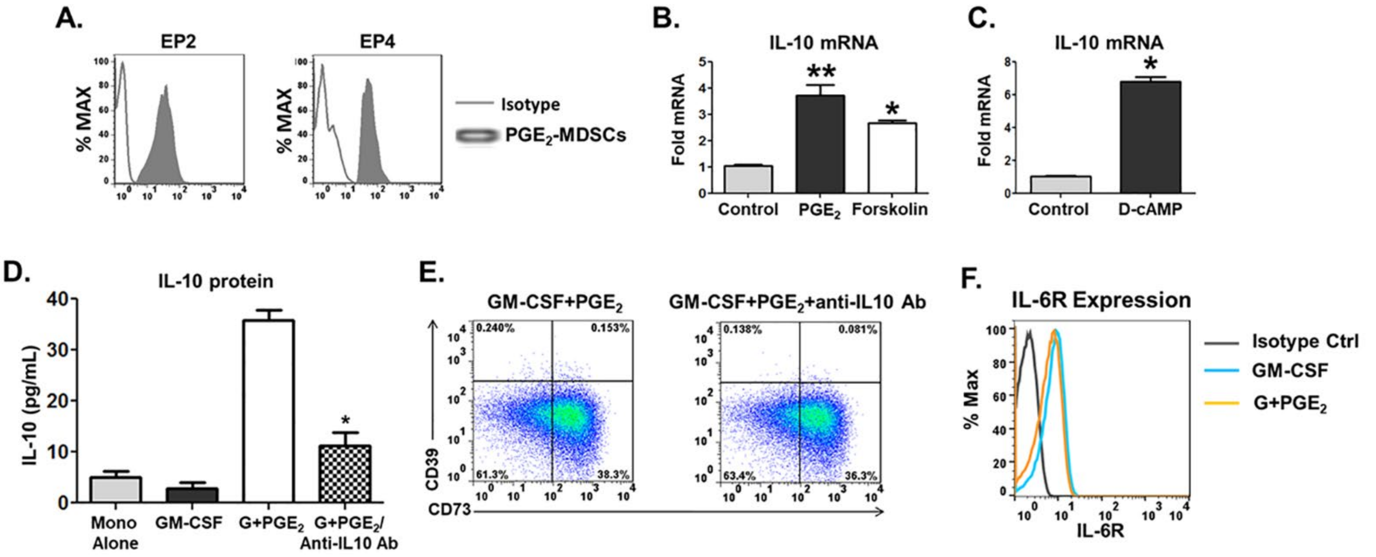


**Supplemental Figure 3. A.** Representative flow cytometry histograms show that $PGE_2$-induced human monocytic MDSCs express both EP2 and EP4 receptors. **B.** Bar graph showing the IL-10 mRNA expression in $CD14^+$ monocytes treated with either vehicle (Control), or $PGE_2$ (1 µM), or Forskolin (10 µM) for 16 hours. Data, mean ± SEM, *P* values: **, p<0.001; *, p<0.01. **C.** Bar graph showing the IL-10 mRNA expression in human $CD14^+$ monocytes treated with either vehicle (Control), or D-cAMP (100 µM) for 16 hours. Data, mean ± SEM, *P* values: *, p<0.01. **D.** Bar graph showing the IL-10 protein expression in untreated human $CD14^+$ monocytes or in monocytes treated with either GM-CSF (10 ng/mL), or GM-CSF and $PGE_2$ (1 µM), or GM-CSF and $PGE_2$/human anti-IL-10 antibody (5 µg/mL) for 16 hours. Data, mean ± SEM, *P* values: *, p<0.01. **E.** Human $CD14^+$ monocytes are treated with either GM-CSF + $PGE_2$/vehicle, or GM-CSF + $PGE_2$/anti-IL-10 antibody for 48 hours, and representative dot plots show the expression of CD39/CD73 in the treated monocytes. **F.** Human $CD14^+$ monocytes treated with either GM-CSF, or GM-CSF and $PGE_2$ and representative flow cytometry histogram shows the surface expression of IL-6 receptor.

### Supplemental Figure 4

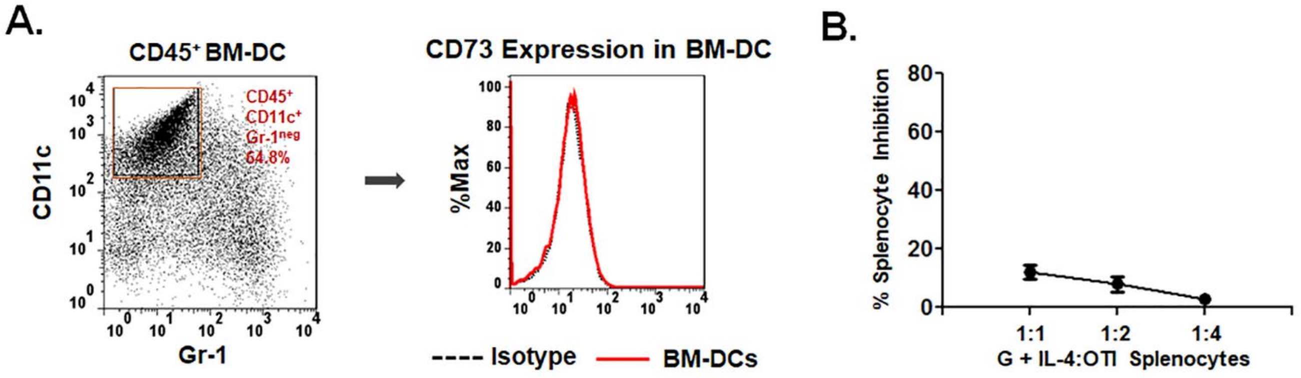


**Supplemental Figure 4. A, B.** Murine bone marrow (BM) cells from naïve C57BL/6 were cultured with either GM-CSF (40 ng/mL) and GM-CSF and IL-4 (10 ng/mL) for 5 days and analyzed by flow cytometry for CD11c, Gr-1, CD39 and CD73 expression and T cell immunosuppressive activity. **A.** Accumulation of $CD45^{+}CD11c^{+}Gr\text{-}1^{neg}$ dendritic cells and expression of CD73 within the $CD45^{+}CD11c^{+}Gr\text{-}1^{neg}$ dendritic cell population are shown in representative histogram. **B.** BM cells were cultured with GM-CSF + IL-4 for 5 days. Cells were harvested and added to splenocytes purified from OT-I transgenic mice at 1:1, 1:2 and 1:4 ratio in the presence of ovalbumin (250 µg/mL) per well for 4 days and T cell proliferation was measured by $^{[3H]}$thymidine incorporation.
